# Control of stress-activated Cdc42 dynamics by the MAP kinase Sty1 – NDR kinase Orb6 regulatory axis

**DOI:** 10.1101/2025.02.26.640195

**Authors:** Laura P. Doyle, Jun-Song Chen, Kathleen L. Gould, Dannel McCollum, Fulvia Verde

## Abstract

Cdc42 is a Rho-family GTPase that controls cell polarization from yeast to human cells. In fission yeast, under normal growth conditions, Cdc42-GTP oscillates between cell tips to promote polarized growth. However, when exposed to environmental stressors, Cdc42 adopts an “exploratory” pattern of Cdc42 activation along the cell membrane. This pattern also occurs when the NDR kinase Orb6 is downregulated. Here, we describe the molecular mechanism behind the emergence of exploratory Cdc42 dynamics and identify a new substrate of Orb6 kinase, the Cdc42 GAP Rga3. Additionally, we show that MAP kinase Sty1, known for linking stress signals to the Cdc42 polarity module, negatively regulates Orb6 kinase. During nutritional stress, activation of Sty1 and inactivation of Orb6 are associated with chronological lifespan extension. Our findings reveal a novel mechanism controlling cell morphology during stress, with important implications for cell survival.

## INTRODUCTION

Cell polarization is an important process involved in cell development, differentiation, migration, and the onset of disease^1–6^. Normal cell polarization is crucial for proper cell movement, asymmetric cell division, and regulation of immune responses^5–11^. Cdc42 GTPase is an essential regulator of polarized cell growth and is evolutionarily conserved in eukaryotes^12,13^. In the yeasts, *S. pombe* and *S. cerevisiae,* Cdc42 plays a key role in establishing polarized cell growth^14–20^. In higher eukaryotes such as *Drosophila melanogaster* and mammalian cells, Cdc42 also plays a central role in cell polarity^21–26^. Cdc42 activity is promoted by guanine-nucleotide exchange factors (GEFs) which facilitate the binding of GTP to Cdc42^27^. Conversely, GTPase activating proteins (GAPs) serve as negative regulators of Cdc42 as they promote the GTPase activity of Cdc42 leading to the hydrolysis of GTP into GDP^27^. We previously showed that in the fission yeast *Schizosaccharomyces pombe*, active Cdc42 oscillates between the tips over time^16,17^. This striking dynamic emerges from positive and negative feedback and competition for Cdc42 regulators and is conserved^16,28–30^. Fission yeast provide an excellent model to study conserved signaling pathways in the control of cell morphogenesis due to its well-defined rod-shaped morphology which allows for straightforward measurement of cell growth and changes in polarization.

Nuclear Dbf2-related (NDR) kinases are a subclass of the AGC (protein A, G, and C) group of serine/threonine protein kinases^31,32^. They are important for various cellular processes involving cell morphogenesis, mitosis, proliferation, and apoptosis and are highly conserved from yeast to humans^31,32^. In higher eukaryotes, NDR kinases have a role in cancer biology, innate immunity, and neuron development^33–38^. In *S. pombe*, the conserved NDR kinase is known as Orb6 and it plays a role in controlling cell polarized growth and cell morphogenesis^16,17,32,39–44^, and it regulates the localization and activity of the key morphological regulator Cdc42 GTPase^16,17,44^.

During interphase and in the presence of nutrients, Orb6 promotes Cdc42 function at the cell tips by increasing the activity of the upstream regulator, Ras1^41^. More specifically, Orb6 phosphorylates the conserved mRNA binding protein Sts5^41,42^. Sts5 phosphorylation increases the protein levels of Efc25, an exchange factor and activator of Ras1 GTPase^41,42^. Ras1 activation recruits the Cdc42 exchange factor Scd1 at the membrane to promote Cdc42 activation and cell polarization during vegetative growth^45,46^. In addition to Sts5, Orb6 kinase also phosphorylates Gef1, the second exchange factor for Cdc42 GTPase, keeping it sequestered in the cytoplasm and inactive^40^. This pattern of Cdc42 activation during vegetative growth when Orb6 is active leads to the anticorrelated Cdc42 oscillations at the cell tips.

Cell polarization is modulated dynamically in response to environmental stresses. Several groups have found that Cdc42 localization changes following activation of the mitogen-activating protein (MAP) kinase, Sty1^41,47–49^, in response to a variety of environmental stressors including osmotic changes, oxidative stress, nutritional starvation, and heat shock^50–55^. Specifically, activation of Sty1 promotes the formation of exploratory patches of active Cdc42 along the lateral cell membrane rather than exclusively at cell tips^47,48,56^. We have previously shown that Orb6 kinase activity decreases upon nutritional stress suggesting that Orb6 downregulation plays a role in cell morphogenesis and cell growth during nutritional deprivation^41^. Consistent with these findings, Orb6 kinase downregulation before stress exposure leads to chronological lifespan extension during nutritional starvation^41^. However, it is currently not known what leads to Orb6 inactivation during nutritional stress or how inactivating Orb6 under these conditions leads to the modulation of active Cdc42 distribution.

In this study, we find that the Cdc42 GTPase activating protein (GAP) Rga3 is a novel substrate of the Orb6 kinase. We show that Orb6 activity negatively regulates Rga3 and limits Rga3 localization to the membrane. Rga3 and the Cdc42 GEF Gef1 cooperate to promote the emergence of exploratory Cdc42 dynamics during nitrogen starvation. Further, we discover that MAP kinase Sty1 activity negatively regulates Orb6 kinase and promotes Gef1 dephosphorylation on the Orb6 site S112. Together, these findings suggest that the switch between polarized growth by the “canonical” Scd1-dependent Cdc42 activation in nutrient-rich conditions to “exploratory” Cdc42 dynamics regulated by Rga3 and Gef1 when Orb6 is inhibited by Sty1 during nutrient stress. Due to the role of Orb6 and Sty1 activity on cell survival, exploratory Cdc42 dynamics during nutrient starvation may have implications in promoting cell resilience during stress.

## RESULTS

### Cdc42 GAP Rga3 membrane localization is regulated by Orb6 kinase activity

We previously showed that Gef1 and Sts5 phosphorylation by Orb6 promotes their binding to 14-3-3 protein Rad24^40–42^. We have found that mutations of these two substrates at their respective Orb6 phosphorylation sites alter cell morphology^40,41^. Mutations of the Orb6 phosphorylation sites in these proteins only partially recapitulate the effects of Orb6 kinase inhibition, however, suggesting that there are other substrates that contribute to the observed changes in Cdc42 dynamics. We reasoned therefore that novel Orb6 substrates might be identified by their association with the 14-3-3 protein, Rad24, in an Orb6-dependent manner. Thus, we used a mass spectrometry screen that was previously employed to successfully identify substrates of the related kinase, Sid2 ^57^. Orb6 and Sid2 belong to the NDR/LATS family of protein kinases that preferentially phosphorylate the consensus sequence [HX(R/K/H)XX(S/T)]^58–63^. When the serine or threonine in this sequence becomes phosphorylated, it forms the consensus binding site for 14-3-3 proteins^40–42,64,65^.

Rad24-TAP complexes were purified from wild-type and temperature-sensitive *orb6-25* cells, both shifted to 36°C for 3 hours. Protein samples were digested and analyzed by two-dimensional liquid chromatography-tandem mass spectrometry (LC-MS/MS) to identify Rad24 binding partners. The abundance (spectral counts) of each Rad24 interactor was normalized to Rad24 abundance. The ratio of individual protein abundance in wild-type and *orb6-25* was compared, revealing several proteins whose abundance declined in the absence of Orb6 function. These were considered putative Orb6 substrates (Table S1). In total, we identified 29 novel candidate substrates of Orb6, 18 of which contain putative Orb6 phosphorylation consensus sequences (Table S2). Validating the approach and supporting our previous findings, Gef1 and Sts5 were identified, which we previously showed are direct substrates of Orb6 kinase and bind to Rad24 in an Orb6-dependent manner^40–42^. The factors identified by mass spectrometry have roles in biological processes involved in cell polarization and morphogenesis including kinase signaling, cytokinesis, metabolic processes, and cell polarity, and many contain consensus sites for Orb6 kinase phosphorylation ([HX(R/K/H)XX(S/T)]^61,66^ (Table S1-S2). In this study, we focused on Rga3, a Cdc42 GAP. Previously we have shown that active Cdc42 oscillates between cell tips when Orb6 is active^16^ (Fig. 1A, Video S1). However, when Orb6 function is inhibited, Cdc42 dynamics change fundamentally. Loss of Orb6 activity leads to dampening of active Cdc42 at the tips and the appearance of ectopic active Cdc42 patches along the lateral cell membrane^44^ (Fig. 1B, Video S2). Rga3 has an established role in promoting exploratory Cdc42 dynamics during mating^67^, but so far was thought to be expendable during vegetative polarized cell growth. To test if Cdc42 dynamics are affected by loss of Rga3 upon Orb6 kinase inhibition, we used analogue-sensitive mutant *orb6-as2* and visualized active Cdc42 (CRIB-GFP) by fluorescent microscopy (Fig. 1C). We found that the loss of *rga3* significantly reduces the formation of active Cdc42 lateral patches upon inhibition of *orb6-as2* mutants with 1-NA-PP1 (Fig. 1D). Furthermore, the amount of active Cdc42 present at the cell tips increased in *rga3Δ* deletion mutants (Fig. 1E). These results demonstrate that the presence of Rga3 is important in promoting exploratory Cdc42 dynamics and reducing Cdc42 at the cell tips upon Orb6 inhibition.

**Figure 1:**
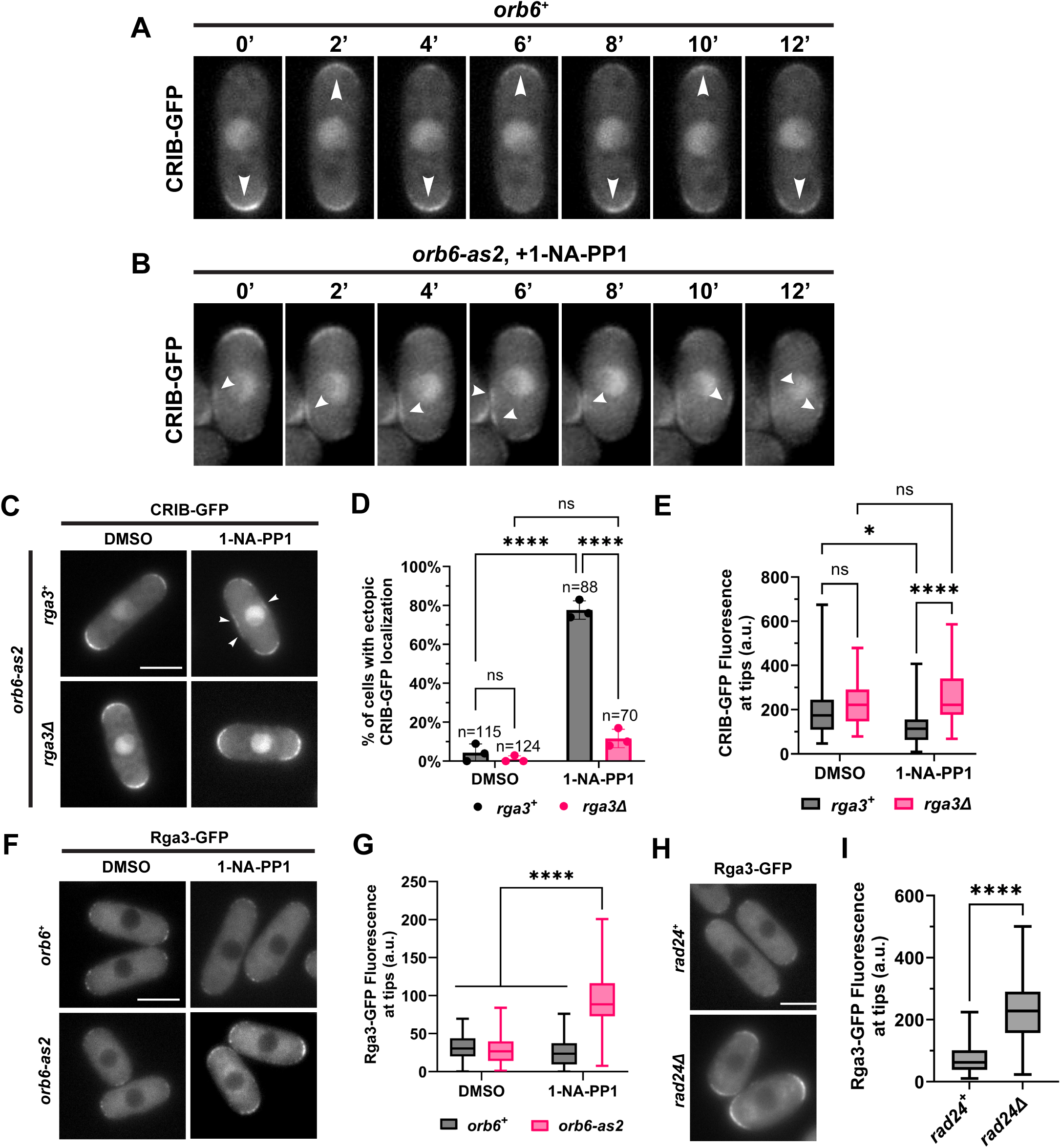
Cdc42 GAP Rga3 membrane localization is regulated by Orb6 kinase activity. **(A)** Active Cdc42 (CRIB-GFP) oscillates between cell tips over time (2-minute intervals). See also Video S1. **(B)** Orb6 inhibition by 1-NA-PP1 leads to the emergence of exploratory active Cdc42 patches along the cell membrane over time (2-minute intervals). See also Video S2. **(C)** Loss of *rga3* reduces the formation of exploratory active Cdc42 (CRIB-GFP) patches along cell membrane upon inhibition of Orb6. Scale bar, 5 μm. **(D)** Quantification of percentage of cells that display ectopic CRIB-GFP localization depicted in C based on three independent experiments. Data presented as mean ± SD, p values are determined by two-way ANOVA with Tukey’s HSD test p ≤ 0.0001, ****. n = number of cells quantified. **(E)** Quantification of CRIB-GFP localization at the cell tips depicted in C based on three independent experiments, n=52 for each group. Whiskers indicating minimum to maximum are shown, box represents 25th to 75th quartiles, and horizontal line represents median, p values determined by two-way ANOVA with Tukey’s HSD test p ≤ 0.05, *; p ≤ 0.0001, ****. **(F)** Rga3-GFP localization at the cell tips increases in *orb6-as2* mutants when Orb6 is inhibited by 1-NA-PP1 for 30 minutes. Scale bar, 5 μm. **(G)** Quantification of F based on three independent experiments. Data presented as in E, p values determined by two-way ANOVA with Tukey’s multiple comparisons test, p ≤ 0.0001,****. **(H)** Rga3-GFP localization at the cell tips increases in *rad24Δ* deletion mutants. Images are a sum projection Z-stack of 6 images separated by a step-size of 0.3 μm. Scale bar, 5 μm. **(I)** Quantification of H, whiskers indicating minimum to maximum are shown, box represents 25^th^ to 75^th^ quartiles, and horizontal line represents median. Welch’s t-test, p ≤ 0.0001,****.

To test if Rga3 localization was affected by Orb6 inhibition, we performed fluorescent microscopy on *wildtype* and *orb6-as2* mutant cells in which Rga3 was tagged with green fluorescent protein (GFP) (Fig. 1F). We found that Rga3-GFP localization significantly increased at the cell tips upon treatment with 1-NA-PP1 in *orb6-as2* cells (Fig. 1G). To corroborate the idea that Rga3 binds to the 14-3-3 protein Rad24, we also visualized Rga3-GFP in the absence of Rad24 and found an increase of Rga3 localization at the cell membrane (Fig. 1H, I). These results indicate that Rga3 localization at the tips is negatively regulated by Orb6 and Rad24 during vegetative polarized growth.

Our screen also identified the polarity factors Tea3 and Tea4, both of which play a role in establishing polarized cell growth^68–70^ as candidate targets of Orb6. Since Tea3 has been reported to physically interact with Rga3 by 2-hybrid^71^, we tested if Orb6 kinase activity affects Tea3 localization and if loss of Tea3 affects the localization of Rga3. We found that, similarly to Rga3, Tea3 localization at the cell tips was increased upon Orb6 inhibition (Fig. S1A, B). Further, Rga3 localization to the cell tips is in part dependent on Tea3 (Fig. S1C, D). Another factor, Tea4, forms a complex with another Orb6 target, the Cdc42 GEF Gef1^72^. Thus, our findings suggest that Orb6 regulates polarity control complexes by promoting the interaction of individual components with Rad24.

### Phosphorylation of serine 683 alters Cdc42 GAP Rga3 localization

There are 4 putative Orb6 phosphorylation sites on Rga3 that share the Orb6 phosphorylation consensus site [HX(R/K/H)XX(S/T)]^61,66^. According to genome-wide phosphoproteomic screens performed by other groups, serine 683 is modified in different conditions including nitrogen starvation and during the cell cycle^66,73^. Thus, to test the role of Orb6 phosphorylation on the S683 site, we converted S683 to alanine using CRISPR/Cas9 to construct a GFP-tagged nonphosphorylatable mutant, *rga3-S683A-GFP,* under the control of the endogenous promoter (Fig. 2A). We found that Rga3-S683A-GFP localization was significantly increased at the tips relative to Rga3-GFP, similar to when Orb6 was inhibited (Fig. 1F and 2B,E). Additionally, we found that Rga3-S683A-GFP localization at the tips did not further increase upon Orb6 inhibition (Fig. S2). This result indicates that loss of S683 phosphorylation is sufficient to explain the Rga3 localization changes observed when Orb6 kinase is inhibited.

**Figure 2:**
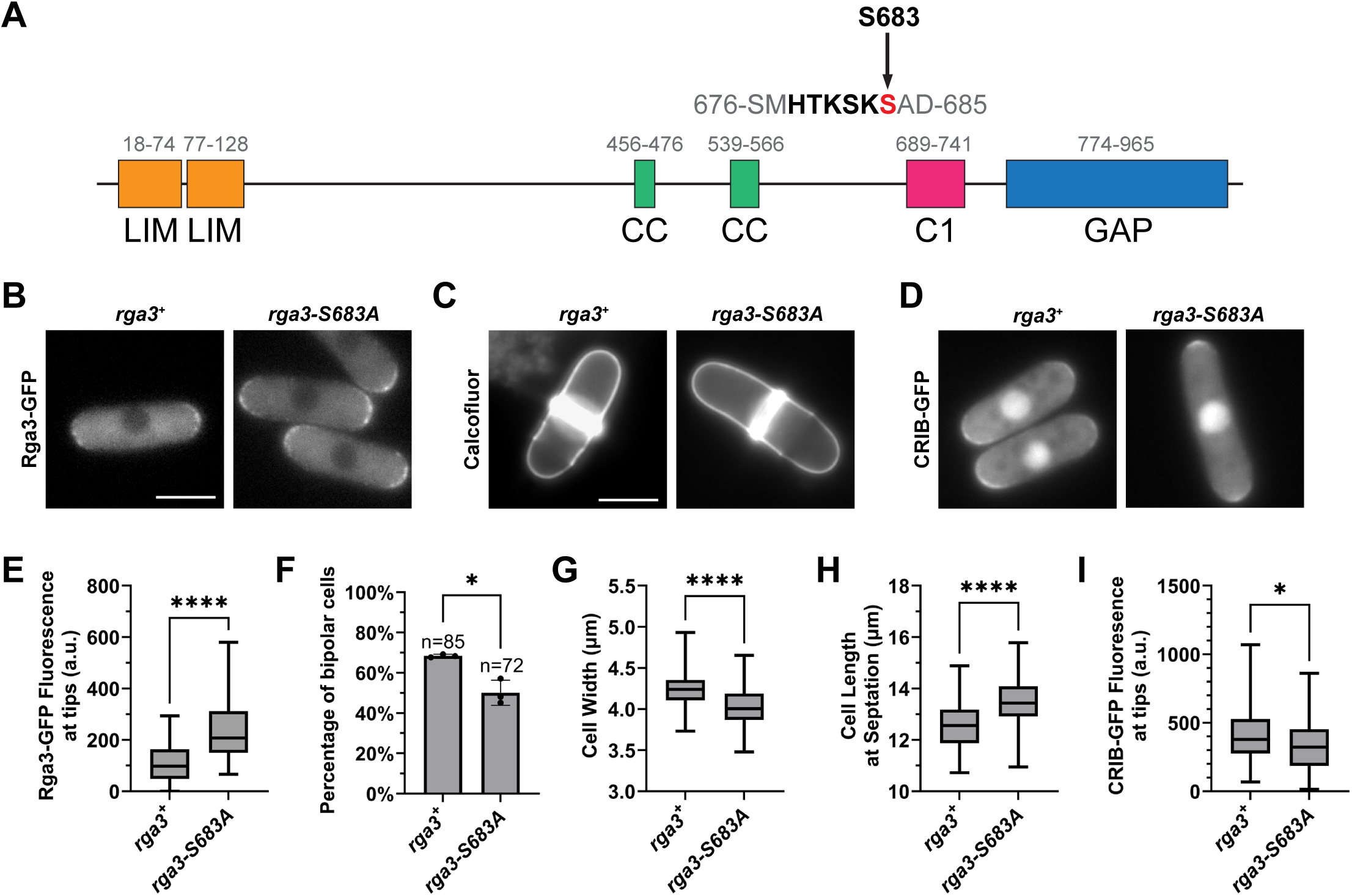
Phosphorylation of serine 683 alters Cdc42 GAP Rga3 localization. **(A)** Protein domain map of Rga3. The arrow indicates serine 683 as the Orb6 consensus phosphorylation site (HX[R/K/H]XX[S/T]). **(B)** Rga3-GFP localization increases in *rga3-S683A-GFP* mutants. Images are a sum projection Z-stack of 6 images separated by a step-size of 0.3 μm. Scale bar, 5 μm. **(C)** Calcofluor staining of *rga3-GFP* and *rga3-S683A-GFP* mutants. Scale bar, 5 μm. **(D)** CRIB-GFP fluorescence at the cell tips decreases in *rga3-S683A-GFP* mutants. Images are a sum projection Z-stack of 6 images separated by a step-size of 0.3 μm. Scale bar, 5 μm. **(E)** Quantification of Rga3-GFP fluorescence depicted in B based on three independent experiments. Whiskers indicating minimum to maximum are shown, box represents 25^th^ to 75^th^ quartiles, and horizontal line represents median, *p* values are determined by Welch’s t-test, p ≤ 0.0001, ****. **(F)** Quantification of the percentage of bipolar cells depicted in D based on three independent experiments. Data are presented as mean ± SD, *p* values are determined by Welch’s t-test, p ≤ 0.05, *. n = number of cells quantified. **(G)** Quantification of cell width depicted in D based on three independent experiments. Data presented as in E, *p* values are determined by two-tailed Student’s t-test, p ≤ 0.0001, ****. **(H)** Quantification of cell length at septation depicted in D based on three independent experiments. Data presented as in E, *p* values are determined by two-tailed Student’s t-test, p ≤ 0.0001, ****. **(I)** Quantification of CRIB-GFP fluorescence at cell tips depicted in H based on three independent experiments. Data presented as in E, *p* values are determined by two-tailed Student’s t-test, p ≤ 0.05, *.

To better understand the role of Rga3-S683 in cell morphology and the control of cell polarization, we imaged cells stained with Calcofluor to visualize the cell wall to measure bipolarity and cellular dimensions (Fig. 2C). As Calcofluor preferentially stains growing cell ends^74^, we can visualize bipolar growth in *S. pombe* through fluorescent imaging of Calcofluor stained cells (detailed in methods). In *rga3-S683A* mutants, we found a decrease in the percentage of bipolar cells (Fig. 2F). Further, we found a significant decrease in cell width and a corresponding increase in cell length of *rga3-S683A* mutant cells (Fig. 2G, H). Decreased percentage of bipolarity and decreased cell width are consistent with increased levels of Rga3 negatively regulating Cdc42 activity at the cell tips^16^. In fact, we observed a decrease in the amount of active Cdc42 present at the tips in *rga3-S683A* cells (Fig. 2D, I). Thus, these results demonstrate that Rga3 localization is altered by phosphorylation at the Orb6 consensus site S683 and that this phosphorylation controls cell morphology and active Cdc42 localization.

### Rga3 and Gef1 promote exploratory Cdc42 dynamics upon attenuation of the vegetative Cdc42 control axis

During nutritional starvation or upon Orb6 kinase inhibition, the Cdc42 GEF Scd1 is reduced at the cell tips in a manner that is mediated by decreased levels of the Ras1 exchange factor Efc25^41^. Concomitantly, active Cdc42 forms dynamic patches along the lateral cell membrane^41,44^. Loss of Gef1 strongly reduces these exploratory Cdc42 patches in *orb6-25* temperature sensitive mutants, indicating that Gef1 is essential for Cdc42 exploratory patch formation^44^. We found that loss of *rga3* significantly also reduced the formation of active exploratory Cdc42 patches upon Orb6 kinase inhibition while simultaneously increasing active Cdc42 at the cell tips (Fig. 1C-E). Therefore, since Orb6 kinase activity decreases during nitrogen starvation, we exposed *rga3Δ, gef1Δ,* and *rga3Δ gef1Δ* deletion mutants to nitrogen starvation to explore a potential cooperative role of Rga3 and Gef1 in controlling Cdc42 dynamics. Here, we found that the double *rga3Δ* and *gef1Δ* deletion mutant maintained the Cdc42 dynamics observed in a non-starved cell (compare Fig. 3Aa and 3Ah), keeping high levels of Cdc42 activity at the cell tips, and abolishing the formation of lateral Cdc42 patches and Cdc42 exploratory dynamics. The single deletion mutant *gef1Δ* almost abolished active Cdc42 from the cell membrane at both the tips and sides, highlighting the critical role of this Cdc42 GEF during nutritional deprivation, and indicating that when Rga3 levels increase at the membrane during nitrogen starvation, Cdc42 activity at the cell tips is repressed (Fig. 3Ag, B, C). Consistent with this idea, the loss of only *rga3Δ* maintained an increased level of active Cdc42 at the tips while significantly reducing the frequency of cells with patches (Fig. 3Af, B, C). These results indicate that both Gef1 and Rga3 cooperate to induce exploratory active Cdc42 dynamics during nutritional deprivation. They also suggest that repression of Cdc42 activity at cell tips is required to promote lateral patch formation.

**Figure 3:**
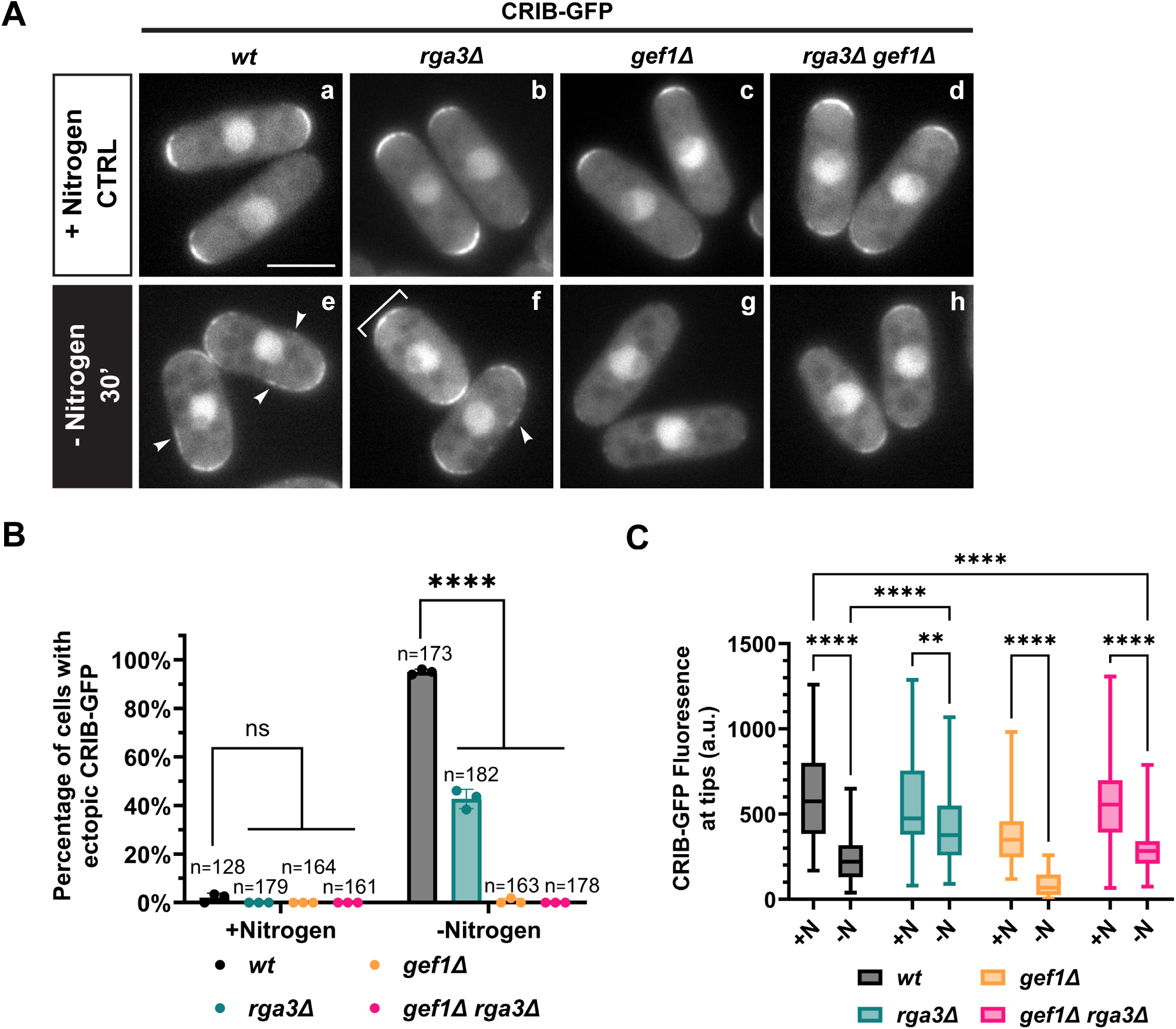
Rga3 cooperates with Gef1 to promote Cdc42 exploratory dynamics during nitrogen starvation. **(A)** Nitrogen starvation leads to the formation of dynamic patches of active Cdc42 along the cell membrane in wild-type cells (*wt,* e). Loss of *gef1* and *rga3* leads to the loss of active Cdc42 patches along the cell membrane and retention of active Cdc42 at the cell tips during nitrogen starvation (h). Scale bar, 5 μm. **(B)** Quantification of percentage of cells that display ectopic CRIB-GFP localization depicted in A based on three independent experiments. Data presented as mean ± SD, p values determined by two-way ANOVA with Tukey’s HSD test p ≤ 0.0001, ****. n = number of cells quantified. **(C)** Quantification of CRIB-GFP localization at the cell tips depicted in A based on three independent experiments. N represents nitrogen. Whiskers indicating minimum to maximum are shown, box represents 25th to 75th quartiles, and horizontal line represents median, p values determined by two-way ANOVA with Tukey’s HSD test p ≤ 0.01, **; p ≤ 0.0001, ****.

Orb6 kinase promotes Ras1 activity and the recruitment of the canonical Cdc42 GEF Scd1 to cell tips by phosphorylation of the mRNA binding protein Sts5^41,45,46^. Upon nitrogen starvation and reduced Orb6 activity, decreased Sts5 phosphorylation leads to a decline in Ras1 activity by lowering Efc25 protein levels^41^. Thus, we investigated how the nonphosphorylatable mutants Rga3-S683A and Gef1-S112A might cooperate when the activity of the primary regulators of Cdc42 during vegetative growth drop, namely in *efc25Δ* deletion mutants, by measuring the frequency of cells displaying ectopic active Cdc42 patches. Remarkably, we found that in *rga3-S683A gef1-S112A efc25Δ* mutants, the frequency of cells displaying ectopic active Cdc42 patches strongly increased, as compared to wild-type, *efc25Δ,* and *efc25Δ* mutants containing either *rga3-S683A* or *gef1-S112A* alone (compare Fig. 4Aa-d and Ae, B).

**Figure 4:**
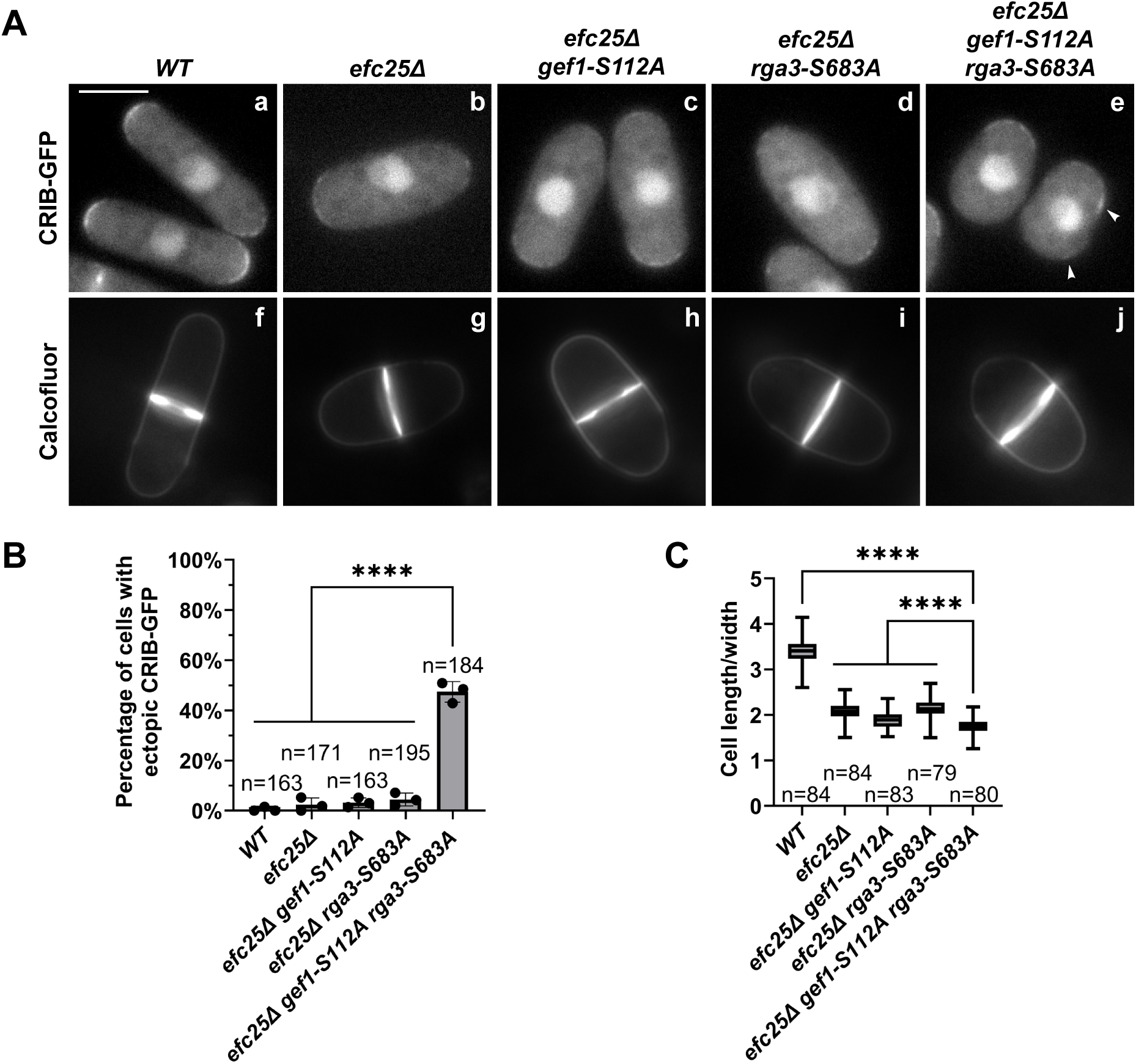
Rga3 and Gef1 promote exploratory Cdc42 dynamics upon attenuation of the vegetative Cdc42 control axis. **(A)** CRIB-GFP fluorescence (a-e) and calcofluor staining (f-j) of *gef1-S112A-HA* and *rga3-S683A* mutants in the background of *efc25Δ*. The triple mutant (*efc25Δ gef1-S112A-HA rga3-S683A*) displays exploratory Cdc42 dynamics and a round phenotype. Scale bar, 5 μm. **(B)** Quantification of percentage of cells that display ectopic CRIB-GFP localization depicted in A,a-e based on three independent experiments. Data presented as mean ± SD, p values are determined by one-way ANOVA with Tukey’s HSD test p ≤ 0.0001, ****. n = number of cells quantified. **(C)** Quantification of the ratio of cell length to cell width at septation depicted in A,f-j based on three independent experiments. Whiskers indicating minimum to maximum are shown, box represents 25th to 75th quartiles, and horizontal line represents median, p values determined by one-way ANOVA with Tukey’s HSD test p ≤ 0.0001, ****. n = number of cells quantified.

To see how these mutants affect cellular dimensions, we measured the ratio of cell length to width of cells undergoing septation. When both *rga3-S683A* and *gef1-S112A* mutations are present in the absence of Efc25, the cells display a rounder phenotype during septation as compared to wild-type (Fig. 4Aj, C). In *efc25Δ* and upon addition of either nonphosphorylatable *gef1-S112A* or *rga3-S683A* mutations, cells undergo septation at a shorter length compared to wild-type (compare Fig. 4Af and Ag-i, C) but are not as round as the triple mutant (Fig. 4Aj, C). These results highlight how preventing Orb6 phosphorylation of Gef1 and Rga3 promotes the formation of active Cdc42 patches when the main regulators of Cdc42 are downregulated. Furthermore, these results demonstrate that Rga3 and Gef1 together are crucial modulators of active Cdc42 dynamics in conditions where Orb6 is inactive, such as during nutritional stress.

### Stress-activated MAP kinase Sty1 negatively regulates Orb6 to promote the emergence of active Cdc42 exploratory dynamics

Although recent research has demonstrated that Sty1 is essential in the emergence of exploratory Cdc42 dynamics during oxidative stress^47^ or actin depolymerization^48^, the mechanism whereby Sty1 controls Cdc42 is still not understood. Here, we explored the effects of Sty1 on Cdc42 dynamics during nitrogen starvation. We found that in the absence of *sty1*, active Cdc42 remained at the cell tips during nitrogen starvation rather than forming patches along the membrane (Fig. 5A, B). Further, while normally Gef1 localizes to the membrane upon nitrogen starvation^41^, we found that in the absence of *sty1*, Gef1 did not localize to the membrane (Fig. 5C, D). Additionally, we found that there was a significant increase in Rga3-GFP localization at the cell tips upon nitrogen starvation and this increase was also dependent on the presence of *sty1* (Fig. 5E, F). Thus, these observations show that Sty1 controls both Gef1 and Rga3 localization to regulate active Cdc42 dynamics during nitrogen stress.

**Figure 5:**
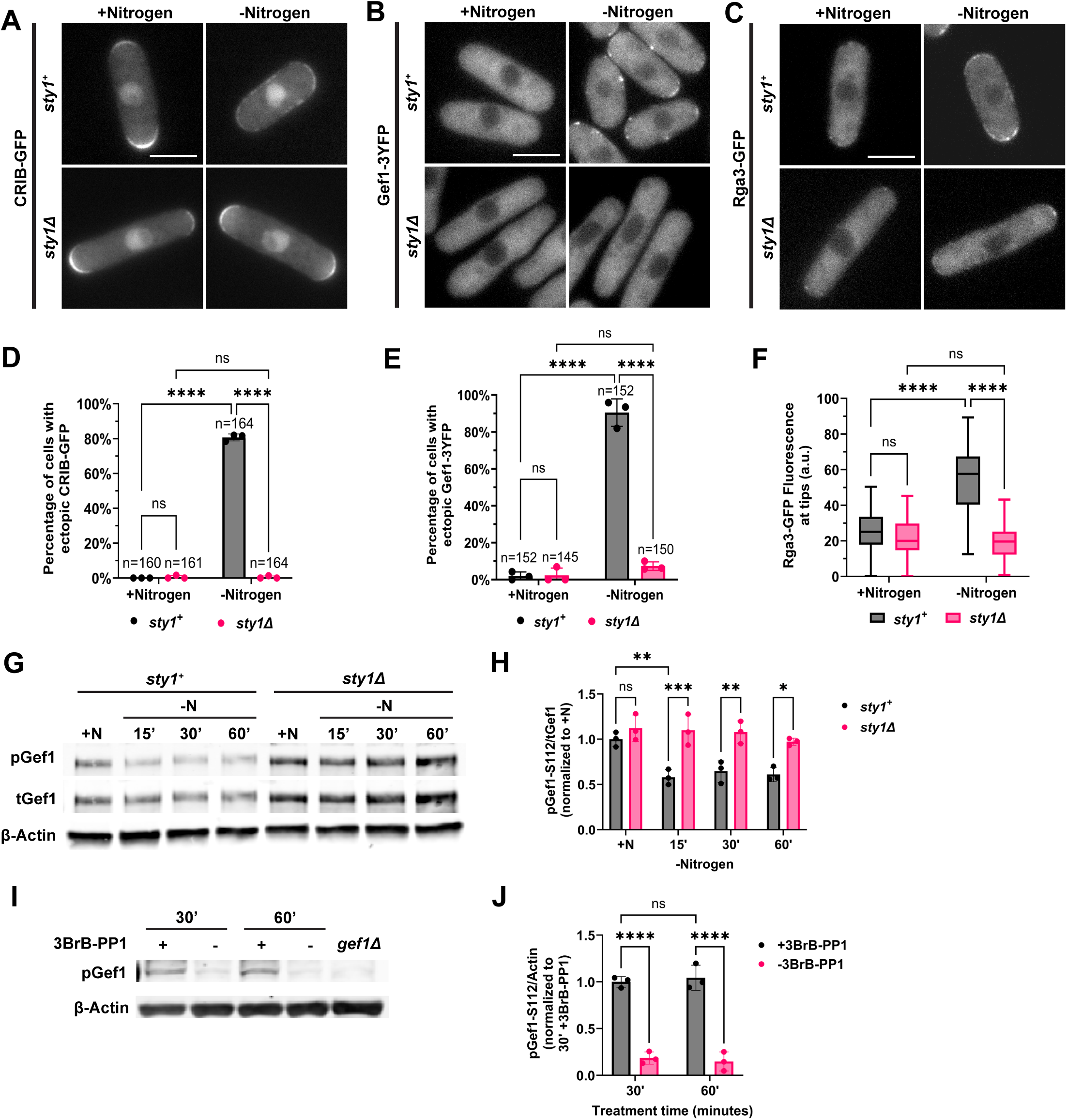
Sty1 activation negatively regulates Orb6 activity. **(A)** Upon nitrogen starvation, active Cdc42 forms patches in *sty1^+^* cells but remains at the cell tips in *sty1Δ* deletion. Scale bar, 5 μm. **(B)** During nitrogen starvation, Gef1-3YFP localizes to the cell membrane but remains sequestered in the cytoplasm when *sty1Δ* is deleted. Scale bar, 5 μm. **(C)** Rga3-GFP is increased at the cell tips upon nitrogen starvation, but Rga3-GFP localization at the tips remains constant in *sty1Δ* deletion cells during nitrogen starvation. Scale bar, 5 μm. **(D)** Quantification of percentage of cells that display ectopic CRIB-GFP localization depicted in A based on three independent experiments. Data are presented as mean ± SD, *p* values are determined by two-way ANOVA with Tukey’s HSD test p ≤ 0.0001, ****. n = number of cells quantified. **(E)** Quantification of percentage of cells that display ectopic Gef1-3YFP localization depicted in B based on three independent experiments. Data presented as in D, *p* values are determined by two-way ANOVA with Tukey’s HSD test p ≤ 0.0001, ****. n = number of cells quantified. **(F)** Quantification of Rga3-GFP localization at the cell tips depicted in C based on three independent experiments. Whiskers indicating minimum to maximum are shown, box represents 25^th^ to 75^th^ quartiles, and horizontal line represents median, *p* values are determined by two-way ANOVA with Tukey’s HSD test p ≤ 0.0001, ****. **(G)** Gef1-S112 phosphorylation by Orb6 remains constant during nitrogen starvation in *sty1Δ* deletion cells. N represents nitrogen. β-Actin was used as a loading control. **(H)** Quantification of pGef1-S112/tGef1 from G during nitrogen starvation in control or *sty1Δ* deletion mutant cells based on three independent experiments. Data presented as in D, *p* values are determined by two-way ANOVA with Tukey’s HSD test p ≤ 0.05, *; p ≤ 0.01, **; p ≤ 0.001, ***. **(I)** Gef1-S112 phosphorylation by Orb6 decreases upon stress-independent activation of Sty1 (−3BrB-PP1). β-Actin was used as a loading control. **(J)** Quantification of pGef1-S112/Actin from I upon Sty1 activation (−3BrB-PP1) based on three independent experiments. Data presented as in D, *p* values are determined by two-way ANOVA with Tukey’s HSD test p ≤ 0.0001, ****.

To test if these effects of Sty1 are mediated at least in part by Orb6 activity, we used a previously described antibody that specifically detects Gef1-S112 phosphorylation by Orb6^41^. In concordance with our previously published results, we found that Gef1-S112 phosphorylation by Orb6 decreases upon nitrogen starvation^41^ (Fig. 5G, H). However, upon the loss of Sty1, Gef1-S112 phosphorylation remained constant during nitrogen starvation (Fig. 5G, H). This result suggests that Sty1 negatively regulates Orb6 kinase activity during stress.

To test if Sty1 kinase activity promotes exploratory active Cdc42 dynamics during nitrogen stress, we used mutants of *wis1,* the MAPKK upstream of Sty1^75,76^. Using CRISPR/Cas9, we created a nonphosphorylatable mutant, *wis1-AA,* and a phosphomimetic mutant, *wis1-DD,* and visualized active Cdc42 upon nitrogen starvation. We found that in the *wis1-AA* mutant, where Sty1 kinase is not phosphorylated by Wis1 and is therefore inactive, Cdc42-GTP remains localized at the cell tips during nitrogen starvation (Fig. S4). These results indicate that Sty1 kinase activity promotes active Cdc42 exploratory dynamics during nitrogen starvation. Additionally, to test if ectopic Sty1 activation inhibits Orb6 kinase in the absence of nitrogen starvation, we measured Gef1-S112 phosphorylation in a strain that allows Sty1 activation without stress stimuli^48^. In this mutant, Sty1 is kept inactive by the presence of the ATP-analog inhibitor 3BrB-PP1 and can be acutely activated by removing the inhibitor. We found that stress-independent activation of Sty1 leads to a significant decrease in Gef1-S112 phosphorylation (Fig. 5I, J). Thus, our results show that Sty1 activation leads to decreased Orb6 kinase activity and that it is Sty1 kinase function that decreases Orb6 activity rather than the stress itself.

To investigate if downregulation of Orb6 kinase activity rescues the phenotypes associated with the loss of *sty1*, we inhibited Orb6 in *orb6-as2* cells with 1-NA-PP1 and measured Gef1-S112 phosphorylation. We found that Gef1-S112 phosphorylation decreased upon Orb6 inhibition irrespective of the presence or absence of *sty1* (Fig. 6A, B), indicating that direct inhibition of Orb6 bypasses a requirement of Sty1 for Gef1-S112 dephosphorylation. Next, we tested the effects of *sty1Δ* on the localization of active Cdc42, Gef1, and Rga3 following inhibition of Orb6. We found that in the absence of *sty1*, inhibition of Orb6 led to exploratory active Cdc42 patch formation at the lateral membrane (Fig. 6C, D). Similar results were obtained when Orb6 was inhibited after nitrogen starvation for 15 minutes (Fig. S3). We found similar localization results for Gef1 (Fig. 6E, F) and Rga3 (Fig. 6G, H). These results show that Orb6 kinase inhibition is sufficient to promote Cdc42 patch formation in the absence of Sty1.

**Figure 6:**
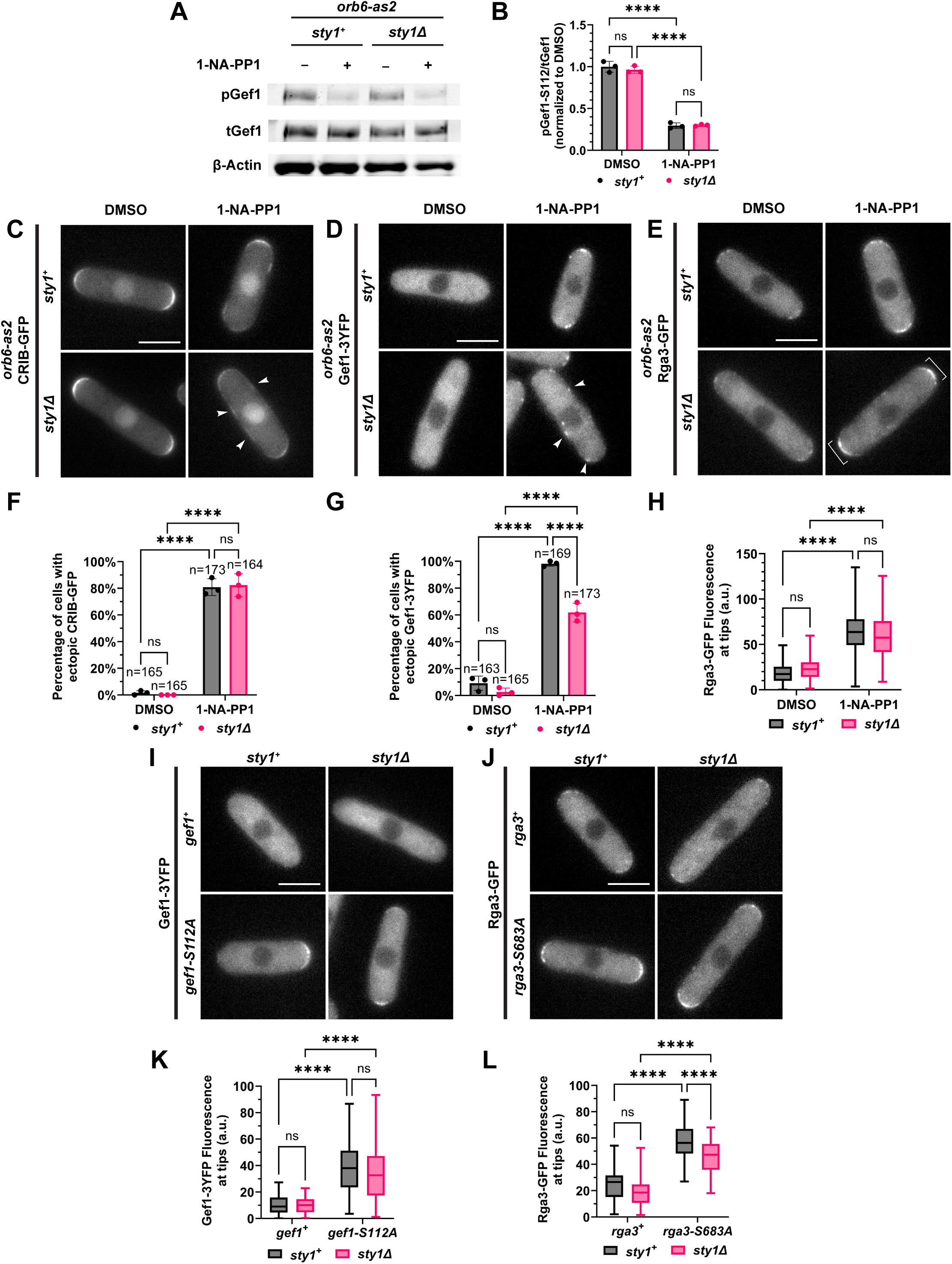
Downregulation of Orb6 kinase activity rescues the phenotypes associated with the loss of *sty1*. **(A)** Gef1-S112 phosphorylation by Orb6 decreases upon Orb6 inhibition (+1-NA-PP1) in *sty1^+^* and *sty1Δ* deletion cells. **(B)** pGef1-S112/tGef1 quantification from A based on three independent experiments. Data are presented as mean ± SD, *p* values determined by two-way ANOVA with Tukey’s HSD test p ≤ 0.0001, ****. **(C)** Inhibition of Orb6 leads to CRIB-GFP patch formation along the cell sides, even in *sty1Δ* deletion cells. Scale bar, 5 μm. **(D)** Gef1-3YFP localizes at the cell membrane during Orb6 inhibition in *sty1Δ* deletion mutants. Scale bar, 5 μm. **(E)** Rga3-GFP increases at the cell tips upon Orb6 inhibition in *sty1Δ* deletion cells. Scale bar, 5 μm. **(F)** Percentage of cells with ectopic CRIB-GFP localization from C based on three independent experiments. Data are presented as in B, *p* values determined by two-way ANOVA with Tukey’s HSD test p ≤ 0.0001, ****. n = number of cells quantified. **(G)** Quantification of percentage of cells that display ectopic Gef1-3YFP localization depicted in E based on three independent experiments. Data are presented as in B, *p* values determined by two-way ANOVA with Tukey’s HSD test p ≤ 0.0001, ****. N = number of cells quantified. **(H)** Rga3-GFP fluorescence quantification at cell tips from G based on three independent experiments. Whiskers indicating minimum to maximum are shown, box represents 25^th^ to 75^th^ quartiles, and horizontal line represents median, *p* values determined by two-way ANOVA with Tukey’s HSD test p ≤ 0.0001, ****. **(I)** Gef1-3YFP localizes to the cell membrane in *sty1^+^* and *sty1Δ* deletion cells in nonphosphorylatable mutant Gef1-S112A-3YFP. Scale bar, 5 μm. **(J)** *sty1Δ* deletion leads to a decrease in Rga3-S683A-GFP localization compared to *sty1^+^.* Scale bar, 5 μm. **(K)** Gef1-3YFP fluorescence quantification at cell tips from I based on three independent experiments. Data presented as in H, *p* values determined by two-way ANOVA with Tukey’s HSD test p ≤ 0.0001, ****. **(L)** Rga3-GFP fluorescence quantification at cell tips from K based on three independent experiments. Data presented as in H, *p* values determined by two-way ANOVA with Tukey’s HSD test p ≤ 0.0001, ****.

Recently, it has been demonstrated that both Gef1 and Rga3 are direct substrates of Sty1^47^. Since Gef1 (as previously shown^40^) and Rga3 (as shown in this study) are also direct substrates of Orb6, we sought to investigate the role of Sty1 in the control of Gef1 and Rga3 localization by Orb6. To do so, we looked at the localization in the nonphosphorylatable mutant of Gef1 or Rga3 by Orb6, Gef1-S112A-3YFP or Rga3-S683A-GFP, in *sty1Δ* cells. Consistent with experiments shown in Fig. 6C-H, we found that Gef1-S112A-3YFP (Fig. 6I, J) and Rga3-S683A-GFP (Fig. 6K, L) localize to the surface independently of Sty1. However, the localization of Rga3-S683A-GFP to the cell tip decreased by about 20% in cells lacking Sty1 indicating that Sty1 promotes Rga3 localization to the cell tips. Similarly, in the absence of Sty1, Gef1 ectopic localization occurred with less frequency, about a 36% decrease, when Orb6 was inhibited, demonstrating that Sty1 plays a role in Gef1 localization to the membrane independently of Orb6 inhibition (Fig. 6E, F). These results demonstrate that loss of Orb6 phosphorylation is the main promoter of Gef1 and Rga3 localization to the membrane. These observations are the first demonstration, to our knowledge, that Sty1 activity also plays a positive role in this effect.

## DISCUSSION

Cdc42, a Rho-family GTPase, has a central role in regulating cell polarization and is highly conserved from yeast to human cells. The ability to control cell polarization is an essential process that changes dynamically in response to environmental stress. In fission yeast, Cdc42 changes the state of polarization in the cell in response to different environmental conditions such as mating^77^, heat shock^49^, oxidative stress^47^, or nitrogen starvation^41^. This change in polarization seen upon exposure to stress is an alternative exploratory pattern of Cdc42 where active Cdc42 localization changes dynamically along the cell membrane rather than oscillating at the cell tips as seen during polarized growth. Our lab has previously shown that inhibition of the nuclear Dbf2-related (NDR) kinase Orb6 activity induces the formation of exploratory active Cdc42 dynamics and extends chronological lifespan^41,44^. We have also shown that exposure to nitrogen starvation leads to a decrease in Orb6 activity^41^. Thus, we sought to understand the molecular mechanisms underpinning the emergence of exploratory Cdc42 dynamics and the role and regulation of Orb6 during stress.

### Exploratory Cdc42 dynamics require dephosphorylation of Cdc42 GAP Rga3 on the Orb6-targeted phospho-site, serine S683

To define novel targets of Orb6 kinase in regulating Cdc42 dynamics, we identified factors that depend on Orb6 kinase for association with 14-3-3 protein Rad24 by mass spectrometry. In this study, we determined that the Cdc42 GAP Rga3 is a novel substrate of Orb6 and found that Orb6 phosphorylation on Rga3 serine 683 limits Rga3 localization to the cell tips. Rga3 is a Cdc42 GAP and plays a contributing role in cell morphology and polarization along with two other Cdc42 GAPs, Rga4 and Rga6^67^. The Rga3 protein structure consists of two LIM domains at the N-terminal, two coiled-coil regions, a C-terminal RhoGAP domain, and is the only Cdc42 GAP that contains a C1 domain in fission yeast^67^. This C1 domain is evolutionarily conserved and thought to have a conserved function in lipid binding from yeast to human^67,78,79^. It has been previously shown that truncating the C1 domain reduces Rga3 localization at the cell tips^67^. In Rga3, serine 683 lies very close to the C1 domain (Fig. 2A), suggesting that Orb6 phosphorylation interferes with C1 domain function by promoting the binding of a bulky 14-3-3 protein, Rad24, and thereby negatively regulating Rga3 localization to the cell membrane. The human orthologs of Rga3 are Rac GAP Chimaerin proteins, CHN1 and CHN2, which also contain C1 domains. Chimaerins have been found to be involved in proper axonal growth and CHN1 mutations can cause a congenital eye movement disorder called Duane Retraction Syndrome^80,81^.

We previously showed that nitrogen starvation decreases Orb6 activity, promoting Cdc42 GEF Gef1 localization to the membrane at ectopic sites^41^. Here, we report that Rga3 plays an essential role in decreasing active Cdc42 at the cells tips and promoting ectopic Cdc42 patch formation upon Orb6 inhibition or during nitrogen starvation. We found that the Orb6 substrates Rga3 and Gef1 cooperate during nitrogen starvation to induce stress-dependent Cdc42 dynamics. When both *rga3* and *gef1* are deleted, Cdc42 remains active at the cell tips, presumably activated by Cdc42 GEF Scd1, and does not form ectopic patches. Deletion of *gef1* alone leads to almost complete abolishment of Cdc42 activation on the whole cell membrane during nitrogen starvation, indicating that increased Rga3 levels at the cell tips during starvation suppress Scd1-dependent Cdc42 activation. Loss of Rga3 alone increases active Cdc42 at the tips and reduces the frequency of cells displaying lateral patches of active Cdc42, during nitrogen starvation, as seen in *rga3Δ* mutants upon Orb6 inhibition. In summary, we conclude that Gef1 and Rga3 cooperate to induce exploratory Cdc42 dynamics in response to nitrogen deprivation. Without Gef1 and Rga3, cells maintain a state of Cdc42 dynamics that is similar to non-starved wild cells, suggesting that Rga3 and Gef1 comprise a nutrient stress-activated module that fosters the emergence of Cdc42 exploratory dynamics in stressed cells. Under nutrient-rich conditions, this Cdc42 regulatory module is kept largely inactive in the cytoplasm by binding to 14-3-3 protein Rad24 (see model in Fig. 7Ba). Decreased Orb6 kinase activity upon nitrogen stress leads to Rga3 and Gef1 dephosphorylation and membrane localization, and activation of exploratory Cdc42 dynamics (Fig. 7, Bb). Rga3 affects the onset of exploratory Cdc42 dynamics also during mating and oxidative stress^47,67^; Gef1 also affects exploratory Cdc42 dynamics during oxidative stress^47^, suggesting that the function of Orb6 in regulating the stress-dependent Cdc42 module may be crucial in the cellular response to other stresses, in addition to nutritional deprivation.

**Figure 7:**
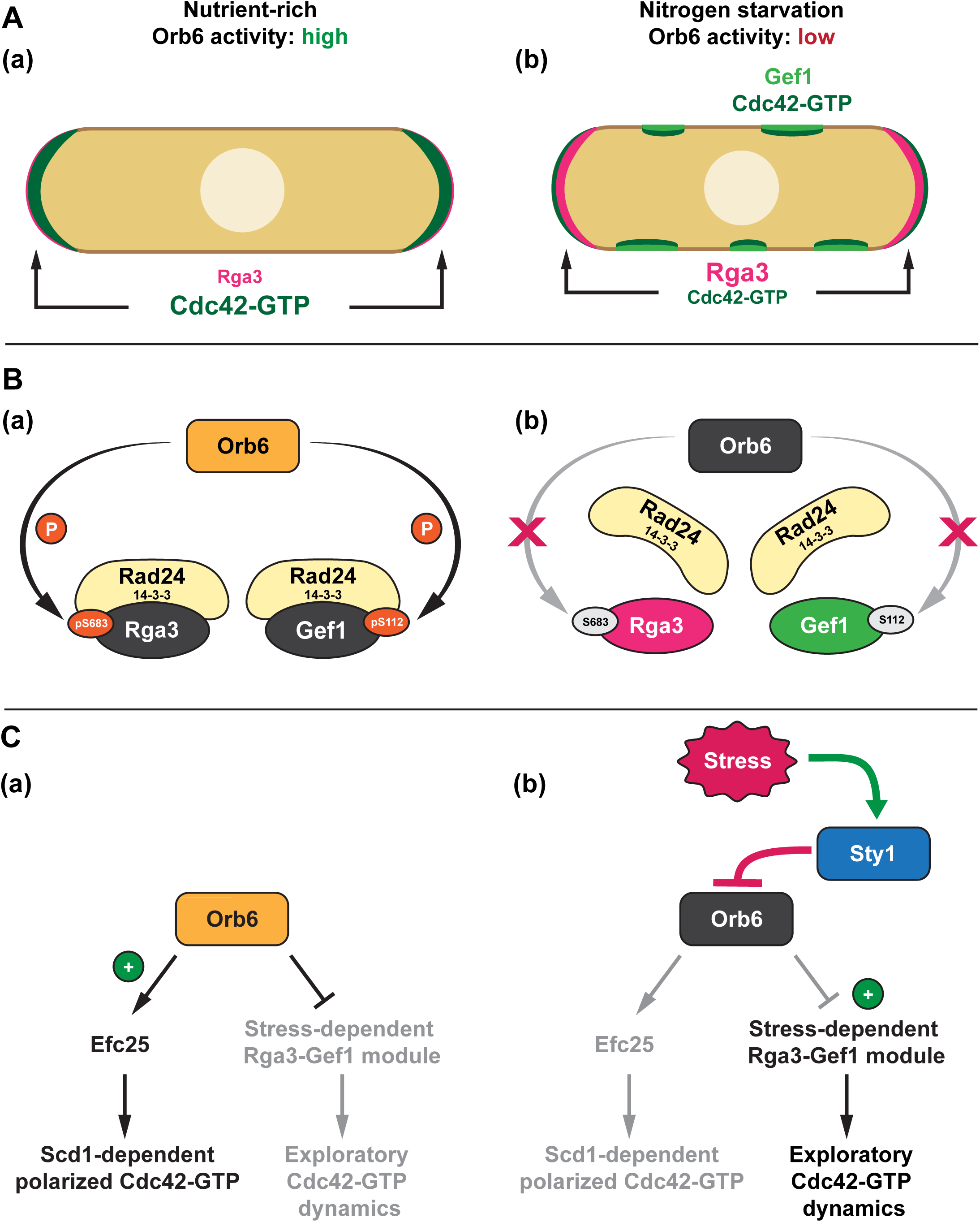
Stress-activated MAP Kinase Sty1 negatively regulates NDR kinase Orb6 to promote exploratory Cdc42 dynamics and cell survival during stress. **(A)** (a) In nutrient-rich conditions, Orb6 remains active promoting active Cdc42 oscillatory dynamics at the cell tips to promote polarized growth. (b) During nitrogen starvation, Orb6 activity decreases leading to the attenuation of Cdc42 activity at the tips and the emergence of exploratory Cdc42 dynamics at the membrane. **(B)** (a) Orb6 phosphorylates Rga3-S683 and Gef1-S112, promoting 14-3-3 Rad24 binding maintaining Rga3 and Gef1 largely sequestered in the cytoplasm. (b) When Orb6 activity decreases during nitrogen starvation, Rga3-S683 and Gef1-S112 become dephosphorylated and are released from Rad24. This release increases Rga3 localization at the cell tips and Gef1 localization to the cell membrane to promote exploratory Cdc42 dynamics. **(C)** (a) Orb6 activation in nutrient-rich conditions stimulates the canonical Cdc42 polarity module to promote polarized growth. (b) Upon stress (nitrogen starvation), Sty1 becomes active and inhibits Orb6 activity. This leads to the induction of the stress-dependent Rga3-Gef1 module to promote exploratory active Cdc42 dynamics.

### The emergence of exploratory Cdc42 dynamics requires the attenuation of the canonical Scd1-dependent Cdc42 polarity module

During nitrogen starvation the canonical Ras1-Scd1-Cdc42 polarity regulatory axis is attenuated^41^. We previously showed that under nutrient rich conditions, Orb6 positively regulates the translation of Ras1 GEF Efc25 (Fig. 7Ca). Upon the onset of nitrogen starvation, or Orb6 inhibition, Ras1 activity and Scd1 localization at the cell tips is attenuated (Fig. 7Cb). We report here that increased levels of Rga3 at the cell tips are crucial to fully repress the canonical Scd1-dependent Cdc42 activity at the cell tips, and thereby promote the alternative exploratory Cdc42 activity. Indeed, we found that in the absence of Efc25, dephosphorylation of Orb6 sites on Gef1-S112 and Rga3-S683 induces the exploratory pattern of Cdc42 activation without stress stimuli. These results together lead us to conclude that upon Orb6 inhibition during nitrogen starvation, Rga3 works in concert with Efc25 downregulation to attenuate active Cdc42 from the cell tips and promote Gef1 at the surface to induce exploratory active Cdc42 dynamics. Here, for the first time to our knowledge, we reveal the molecular mechanism behind the emergence of exploratory Cdc42 dynamics seen during stress.

### MAP kinase Sty1 activity negatively regulates Orb6 kinase to induce exploratory Cdc42 dynamics

One crucial question pertains to how Orb6 kinase activity is downregulated during nutritional stress. Here, we report the novel finding that the stress-activated mitogen-activated protein (MAP) kinase, Sty1, promotes Orb6 inactivation during nitrogen starvation. Sty1 is activated upon exposure to various types of stressors including heat stress, oxidative stress, osmotic changes, and nutritional starvation^50–55^. Sty1 has been shown to alter active Cdc42 localization and dynamics during oxidative stress ^47^. Here we report that also during nitrogen starvation, exploratory Cdc42 dynamics depend on the presence of Sty1. Consistent with this finding, Gef1 and Rga3 localization to the cell membrane is also Sty1-dependent, explaining why active Cdc42 does not form dynamic ectopic patches during nitrogen starvation in *sty1* deletion mutants.

Consistent with the idea that Sty1 functions upstream of Orb6 kinase, we find that phosphorylation of Gef1 on Serine S112, a site which is specifically phosphorylated by Orb6 kinase^40^ remains high in *sty1* deletion cells during nitrogen starvation, indicating that Gef1-S112 dephosphorylation is dependent on Sty1. Conversely, ectopic Sty1 activation in the absence of stress leads to Gef1-S112 dephosphorylation, indicating that Orb6 kinase activity decreases upon Sty1 activation. Remarkably, inhibition of Orb6 completely bypasses the requirement of Sty1, triggering the emergence of exploratory active Cdc42 dynamics, Gef1 lateral patch formation, and increased Rga3 localization at the cell tips. Thus, we show that Orb6 inhibition is necessary and sufficient to induce active Cdc42 exploratory dynamics downstream of Sty1 kinase.

Previous reports showed that Sty1 activation, either stress-independent or by Latrunculin A, induces lateral patches of active Cdc42 on the cell membrane^48^. These results identified Sty1 as a regulator of active Cdc42 localization but did not expand on the mechanism behind Sty1 control of Cdc42 dynamics. Elegant work from the Hidalgo lab showed that Sty1 activation during oxidative stress promotes exploratory Cdc42 dynamics and that Sty1 directly phosphorylates Gef1 and Rga3 *in vitro*, identifying these two proteins as stress-dependent Sty1 targets in the control of Cdc42 dynamics^47^. However, while they identified 7 possible Sty1 phosphorylation sites on Gef1 from amino acids 1-307 and 17 possible sites on Rga3 from amino acids 1-447, they could not report any phenotype associated with phosphomimetic or hypophosphorylated mutants of Gef1^47^. Indeed, this could be explained by the fact that Orb6 inhibition is a crucial step downstream of Sty1 activation. Orb6 inactivation leads to the release of Rga3 and Gef1 from the grip of Rad24, allowing then direct Sty1 phosphorylation of Rga3 and Gef1 to perform a more subtle role in modulating Gef1 and Rga3 membrane localization. In the absence of Orb6 inactivation, Gef1 and Rga3 stay cytoplasmic, and the effects of mutations in the Sty1 consensus sites would likely be muted. Additionally, other factors that are in a protein complex with Gef1 or Rga3 may also need to be dephosphorylated: for example, Gef1 is recruited by Tea4-PP1 to activate Cdc42 at the cell membrane^72^. We found that Tea4 also interacts with Rad24 in an Orb6-dependent manner. Thus, our data is consistent with a stepwise model, where, during stress (1) Sty1 becomes activated and inhibits Orb6, (2) Rga3 and Gef1 are dephosphorylated at their Orb6 site, released by 14-3-3 Rad24, and localize to the membrane, and (3) Sty1 phosphorylation further promotes Rga3 and Gef1 function. This mechanism can be described as type of type 4 circuitry called a coherent feedforward loop^82^, where a double inhibition (Sty1 inhibiting Orb6, Orb6 inhibiting Gef1 and Rga3) occurs in parallel with activation (Sty1 subtly promoting Gef1 and Rga3 localization, and Rga3 GAP activity^47^).

It should also be noted that Orb6 inactivation downstream of Sty1 activation also provides an explanation for the attenuation of the canonical, Scd1 dependent Cdc42 activity observed during oxidative stress^41,47^. We previously showed that Orb6 inactivation leads to the downregulation of Efc25-Ras1 regulatory axis, leading to decreased recruitment of the Cdc42 GEF Scd1 at the tips^41^. Here we show that Orb6 inactivation leads to increased levels of the Cdc42 GAP Rga3 at the tips. Although the data was not shown, the Hidalgo lab has reported that Sty1 phosphorylation leads to the activation of Rga3 GAP activity to promote the hydrolysis of Cdc42-GTP^47^. Thus, these observations collectively support the idea that Orb6 inactivation and Sty1 activation cooperate to promote Rga3 function, resulting in additional suppression of the Cdc42 activity at the tips. It appears that silencing the canonical Cdc42 pathway at the cell tips is a crucial step to allow the emergence of Cdc42 exploratory dynamics.

During nutritional deprivation, cells must be able to adapt to the environment appropriately to promote cell resilience and survival^83^. Chronic nutrient starvation can lead the cells to enter a state of quiescence, a reversible process that occurs when cell division stops and leads to improved cell survival until nutrients become available (reviewed by Yanagiba^84^ and Valcourt et al.^85^). It has been found that Sty1 kinase is activated during stationary phase, when the cells are quiescent^86^. Sty1 and other conserved signaling kinases such as TOR kinase are important regulators of chronological lifespan^87–90^. Chronological lifespan assays, performed under nutrient restriction, provide a readout of cell resilience to stress^91–93^. Sty1 activation is crucial in extending chronological lifespan during caloric restriction^87^ and TORC1 inhibition by caffeine and rapamycin also leads to lifespan extension^88^. Previously, we have shown that Orb6 activity decreases when cells enter a state of quiescence^41^. We have also shown that downregulation of Orb6 prior to entering cell quiescence leads to chronological lifespan extension^41^. The mRNA binding protein Sts5 is an Orb6 substrate that is important in extending chronological lifespan^41^. Orb6 phosphorylation of Sts5 promotes the translation of many polarity factors, including Efc25, a GEF of Ras1 GTPase. Inactivation of Orb6 during nutritional deprivation promotes the downregulation of Ras1 activity, a key event in extending cell survival^41^. Ectopic Ras1 activation shortens, while Ras1 activity downregulation extends chronological lifespan^41^, suggesting that continued activity of the canonical Cdc42 polarity complex at the cell tips is detrimental during starvation. Cdc42 has a known function in polarized secretion and cell wall biogenesis^94,95^ and may be crucial for the necessary changes in the cytoskeleton and cell wall remodeling during stress. Indeed, preliminary experiments suggest that the loss of both *rga3* and *gef1* is shorter lived than controls suggesting that decreasing exploratory active Cdc42 dynamics during starvation is detrimental to the cell (data not shown). These findings coupled with the results of this study suggest an important function for exploratory Cdc42 dynamics upon negative regulation of Orb6 by Sty1 in promoting cell survival and cell adaptation to stress. Further research remains to be performed to elucidate how these Cdc42 dynamics could improve cell resilience during stress. Interestingly, in mice and human cells, increased levels of active Cdc42 and “apolar distribution” of active Cdc42 are detected upon aging^96,97^, highlighting the conserved role of cell polarity control factors in this process.

In conclusion, here we demonstrate that MAP kinase Sty1 is a negative regulator of Orb6 kinase during nutritional stress, promoting the onset of exploratory Cdc42 dynamics. In this proposed model (Fig. 7), Sty1 is activated upon stress and negatively regulates Orb6 inducing the exploratory pattern of Cdc42 dynamics mediated by Gef1 and Rga3 (Fig. 7Cb). Our findings reveal the elegant mechanism whereby Sty1 and Orb6 kinases control the emergence of an alternative pattern of Cdc42 dynamics during stress. The interplay between Orb6 inactivation and Sty1 activation allows the transition from a “canonical” to an “exploratory” active Cdc42 distribution. Since NDR kinases and stress-activated MAP kinases are highly conserved from yeast to human, both structurally and functionally, uncovering the mechanisms regulating stress signaling and cell polarization could provide valuable insight into human health and onset of disease.

### Limitations of the study

In this study, we find that Sty1 negatively regulates Orb6 activity to induce an alternative exploratory pattern of active Cdc42 dynamics during stress. Whether Sty1 is directly inhibiting Orb6 remains to be elucidated, although Orb6 contains several putative Sty1 phosphorylation sites (S/TP) which could mediate this response. Other components of Orb6 signaling module could also be involved such as Mob2 and Nak1, which contain one and ten putative Sty1 phosphorylation sites, respectively. It is important to acknowledge that nitrogen starvation triggers many different pathways of the cell besides Orb6 and Sty1, perhaps most notably, the growth and stress control complexes TORC1/2. As TORC1/2 play key roles in cell growth and nutrient sensing and Sty1 has been found to interact with Sin1, a part of the TORC2 complex, further studies will need to be performed to elucidate the connection between Orb6, Sty1, and TORC1/2 and how they may regulate stress signaling^98–103^.

## RESOURCE AVAILABILITY

### Lead contact

Further information and requests for resources should be directed to the lead contact, Fulvia Verde.

### Materials availability

All unique and stable reagents generated in this study are available from the lead contact.

### Data and code availability

- Data reported in this paper is available upon request.
- This paper does not report any original code.
- Any additional information required to reanalyze the data reported in this paper is available from the lead contact upon request.

## ACKNOWLEDGMENTS

We thank Sophie Martin (University of Geneva, Geneva, Switzerland) and James Moseley (Geisel School of Medicine at Dartmouth, Hanover, New Hampshire, USA) for sharing strains. We thank Daniel Isom for his helpful guidance on using CRISPR/Cas9 and Vera Mariani for assisting in data acquisition. This work was supported by the National Institute of Health (NIH) R01 GM129514 and by the Sylvester Comprehensive Cancer Center (PG012722) at the University of Miami.

## AUTHOR CONTRIBUTIONS

Conceptualization, L.D., D.M., K.G., and F.V.; Methodology, L.D. and F.V.; Investigation, L.D., J-S.C., D.M., and F.V.; Writing - L.D., J.-S.C., K.G., D.M., and F.V.

## STAR★METHODS

### KEY RESOURCES TABLE

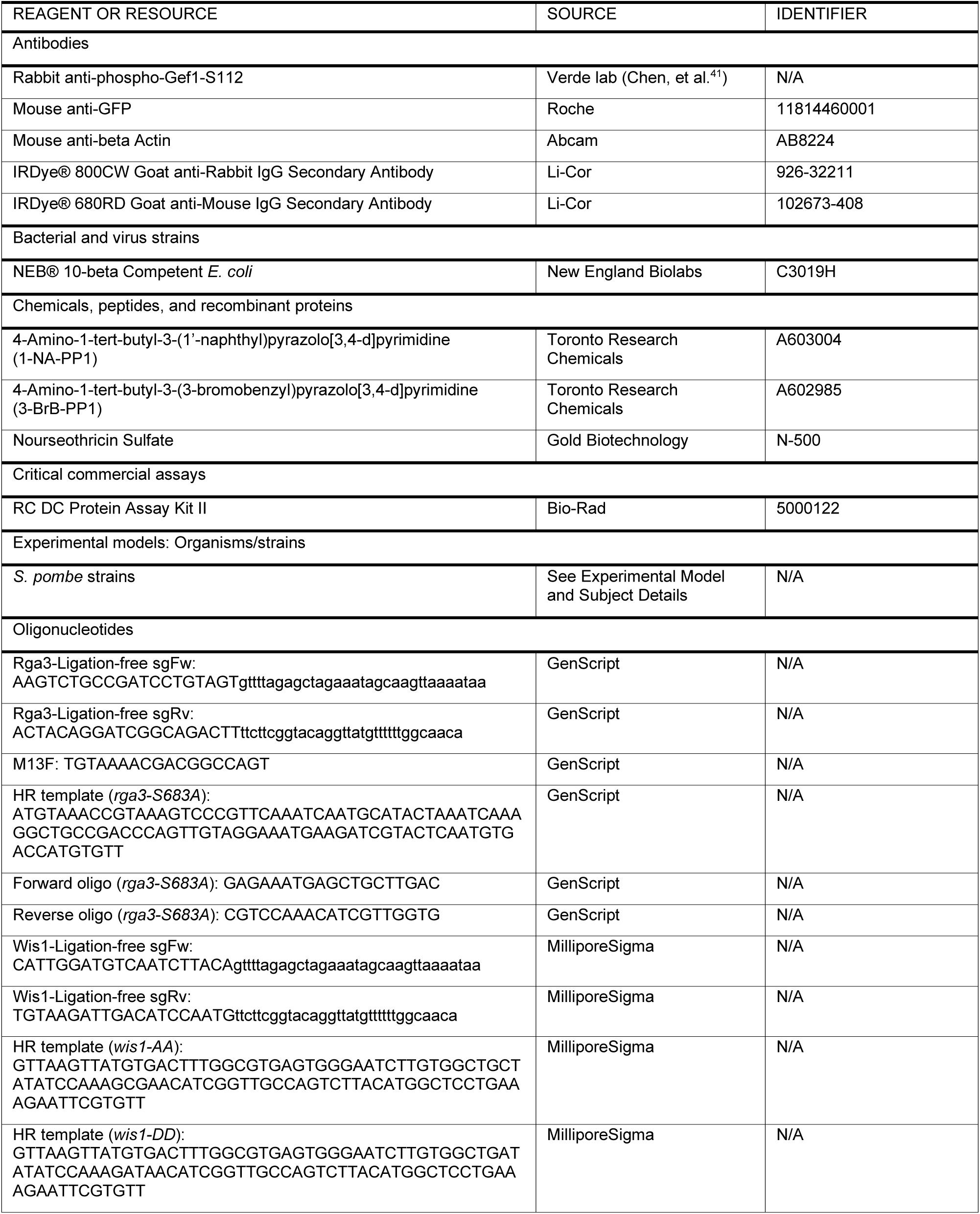
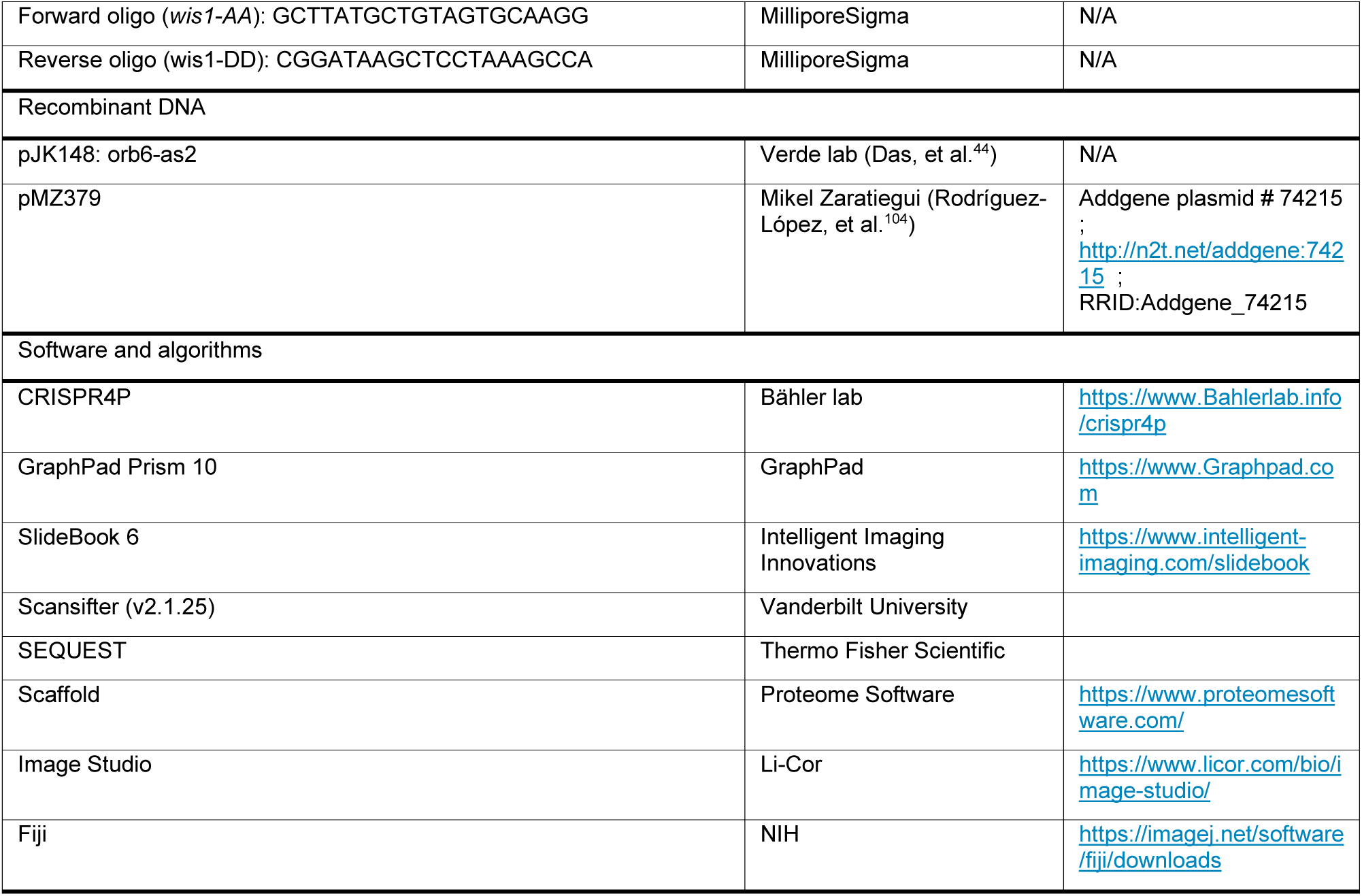

### EXPERIMENTAL MODEL AND STUDY PARTICIPANT DETAILS

*S. pombe* strains used in this study are listed below. All strains used in this study are isogenic to the original strain 972. Fission yeast cells were grown in a shaking incubator at 180 RPM at 25°C and cultured in Edinburgh minimal medium (EMM) plus required supplements unless otherwise specified. Cells used in nitrogen starvation experiments were prototrophic and were cultured in unsupplemented EMM +/− 0.5% nitrogen resource (NH_4_Cl) and grown at 30°C. Experiments using strains with *sty1Δ* deletion or *wis1* mutants were grown at 30°C. Exponential growth was maintained for at least eight generations before experiments, and genetic manipulations and analyses were carried out following standard techniques^105^.

| Strain | Genotype | Origin |
| --- | --- | --- |
| 972 | *h-* | P. Nurse |
| FV3186 | *h- rga3-GFP::kanMX6 ade6- leu1-32 ura4-D18* | This study, <sup>67</sup> |
| FV3166 | *rga3-GFP::kanMX6 orb6Δ::ura+ pJK148orb6-as2::leu+ ade-* | This study |
| FV3539 | *rad24Δ::kanMX6 rga3-GFP::kanMX6 ade6- leu1-32 ura4-D18* | This study |
| FV3409 | *h- rga3-S683A-GFP::kanMX6 ade6- leu1-32 ura4-D18* | This study |
| FV2290 | *h- CRIB-GFP::ura+ leu1-32* | <sup>27</sup> |
| FV3417 | *rga3-S683A CRIB-GFP::ura+ ade6- leu1-32* | This study |
| FV1495 | *gef1Δ::ura4+ gef1-HA::kanMX6 CRIB-GFP::ura+ leu1-32 ade-* | <sup>44</sup> |
| FV3555 | *efc25Δ::kanMX6 gef1Δ::ura4+ gef1-HA::kanMX6 CRIB-GFP::ura+ leu1-32 ade-* | This study |
| FV3496 | *efc25Δ::kanMX6 gef1Δ::ura4+ gef1-S112A-HA::kanMX6 CRIB-GFP::ura+ leu1-32 ade-* | This study |
| FV3557 | *efc25Δ::kanMX6 gef1Δ::ura4+ gef1-HA::kanMX6 rga3-S683A CRIB-GFP::ura+ leu1-32 ade-* | This study |
| FV3498 | *efc25Δ::kanMX6 gef1Δ::ura4+ gef1-S112A-HA::kanMX6 rga3-S683A CRIB-GFP::ura+ leu1-32 ade-* | This study |
| FV1109 | *orb6Δ::ura+ pJK148orb6-as2::leu+ ade- CRIB-GFP::ura+* | <sup>44</sup> |
| FV3408 | *rga3Δ::kanMX6 orb6Δ::ura+ pJK148orb6-as2::leu+ ade- CRIB-GFP::ura+* | This study |
| FV3523 | *gef1Δ::ura+ CRIB-GFP::ura+ ade+ leu+* | This study |
| FV3522 | *rga3Δ::kanMX6 CRIB-GFP::ura+ ade+ leu+* | This study |
| FV3528 | *rga3Δ::kanMX6 gef1Δ::ura+ CRIB-GFP::ura+ ade+ leu+* | This study |
| FV3527 | *CRIB-GFP::ura+ ade+ leu+* | <sup>41</sup>, this study |
| FV3712 | *CRIB-GFP::ura+ sty1Δ::kanMX6 ade+ leu+* | This study |
| FV2206 | *gef1Δ::natMX6 gef1-3YFP::kanMX6 ade+ leu+ ura+* | <sup>41</sup> |
| FV3634 | *gef1Δ::ura+ gef1-3YFP::kanMX6 sty1Δ::kanMX6 ade+ leu+* | This study |
| FV3476 | *rga3-GFP::kanMX6 ade+ leu+ ura+* | This study |
| FV3732 | *rga3-GFP::kanMX6 sty1Δ::kanMX6 ade+ leu+ ura+* | This study |
| FV3702 | *rga3-S683A-GFP::kanMX6 sty1Δ::kanMX6 ade6- leu1-32 ura4-D18* | This study |
| FV1475 | *gef1Δ::ura+ gef1-S112A-3YFP::kanMX6 ade- leu1-32* | <sup>40</sup> |
| FV3697 | *gef1Δ::ura+ gef1-S112A-3YFP::kanMX6 sty1Δ::kanMX6 ade- leu1-32* | This study |
| FV1109 | *orb6Δ::ura+ pjkor6-as2::leu+ CRIB-GFP::ura+ ade-* | <sup>44</sup> |
| FV3734 | *sty1Δ::kanMX6 orb6Δ::ura+ pjkor6-as2::leu+ CRIB-GFP::ura+ ade-* | This study |
| FV3686 | *gef1-3YFP::kanmx orb6Δ::ura+ pjkor6-as2::leu+ ade+* | This study |
| FV3689 | *sty1Δ::kanmx gef1-3YFP::kanmx orb6Δ::ura+ pjkor6-as2::leu+ ade+* | This study |
| FV3719 | *sty1Δ::kanmx rga3-GFP::kanmx orb6Δ::ura+ pjkor6-as2::leu+ ade-* | This study |
| JM4240 | *h+ sty1-T97A wis1DD:12myc::ura4+ pyp1Δ::ura+ pyp2Δ::LEU2 ars1(Blp1):Padh13:CRIB-3xmCitrine:LEU2 leu1-32 ura4-D18* | <sup>106</sup>, <sup>48</sup> |
| YDM4812 | *h- orb6-25 rad24-3HA-TAP::ura+* | D. McCollum laboratory |
| YDM4600 | *h- pj210 rad24-3HA-TAP::ura4+* | D. McCollum laboratory |
| FV3749 | *CRIB-GFP::ura+ wis1-AA ade+ leu+* | This study |
| FV3750 | *CRIB-GFP::ura+ wis1-DD ade+ leu+* | This study |

### METHOD DETAILS

#### Rad24-TAP purification

Rad24-3HA-TAP was purified from *wildtype* and *orb6-25* cells as described previously^107^. Cells were grown in 1.5-liter cultures of 4x YE media at 25°C to 1.6 OD, shifted to 35.5 °C for 3 h., collected by centrifugation at 3000 RPM, and frozen in liquid nitrogen. Cells pellets were thawed on ice and washed with NP-40 Buffer (1% NP-40, 150 mM NaCl, 2 mM EDTA, 6 mM Na2HPO4, and 4 mM NaH2 PO4) supplemented with yeast protease inhibitor cocktail (Sigma-Aldrich), 1 mM phenylmethylsulfonyl fluoride (PMSF), and phosphatase inhibitors (50 mM NaF, and 100 uM NaVO4). Cells were lysed on ice using a bead beater (Biospec) filled with approximately 250 ml of glass beads (425–600 µm G8772 Sigma). The bead beater chamber was immersed in an ice-water mix and run for 8 cycles of 30 s with 30 s cooling periods in between each cycle. Beads were then extracted three times with 50 ml of NP-40 buffer with inhibitors (see above). Cells extracts were then cleared by centrifugation at 3000 RPM for 5 min at 4°C. Subsequent TAP purification was performed as described^107^.

#### Analysis of TAP Complexes by Mass Spectrometry (MS)

Proteins were digested by trypsin and analyzed by two-dimensional liquid chromatography tandem mass spectrometry (MS) (2D-LC-MS/MS) as previously described^108^. MS2 and MS3 spectra were extracted separately from RAW files and converted to DTA files using Scansifter software^109^ (v2.1.25). Spectra with less than 20 peaks were excluded and the remaining spectra were searched using the SEQUEST algorithm (ThermoFisher Scientific, San Jose, CA, USA; version 27, rev. 12). Sequest was set up to search the S. pombe protein database (pombe_contams_20151012_rev database, created in October 2015 from pombase.org). Common contaminants were added, and all sequences were reversed to estimate the false discovery rate (FDR), yielding 10390 total entries. Variable modifications (C+57, M+16, [STY]+80 for all spectra and [ST]-18 for MS3), strict trypsin cleavage, <10 missed cleavages, fragment mass tolerance: 0.00 Da (because of rounding in SEQUEST, this results in 0.5 Da tolerance), and parent mass tolerance: 2.5 Da were allowed. Peptide identifications were assembled and filtered in Scaffold (v4.8.4, Proteome Software, Portland, OR) using the following criteria: minimum of 99.0% protein identification probability; minimum of two unique peptides; minimum of 95% peptide identification probability. FDRs were estimated in Scaffold based on the percentage of decoy sequences identified after using the above filtering criteria; for the combined MudPITs, the protein level FDR was 0.4% and the peptide level FDR was 0.1%. Proteins containing the same or similar peptides that could not be differentiated based on MS/MS alone were grouped to satisfy the principles of parsimony.

#### Fluorescence microscopy

Cells expressing fluorescently tagged proteins were photographed with an Olympus fluorescence BX61 microscope (Melville, NY) equipped with Nomarski differential interference contrast (DIC) optics, a 100× objective (numerical aperture [NA] 1.35), a Roper Cool-SNAP HQ camera (Tucson, AZ), Sutter Lambda 10 + 2 automated excitation and emission filter wheels (Novato, CA), and a 175-W Xenon remote source lamp with liquid light guide. Images were acquired and processed using Intelligent Imaging Innovations SlideBook image analysis software (Version 6.0.4; Denver, CO) and prepared with Fiji^110^. For the measurements of tip Rga3-GFP and CRIB-GFP intensity, we measured the intensity of fluorescence at the tips and subtracted the cytoplasmic background for each cell. 20 cells (40 cell tips) were measured per experiment totaling to 60 cells (120 cell tips) for 3 experiments, unless otherwise specified in which case the number of cells quantified (n=) is included on the figure. To measure cell length and width, cells were stained with Calcofluor at a working concentration of 2μg/mL and Fiji line tool was used to measure length and width of cells undergoing septation. As calcofluor stains growing cell ends brighter, we measured the percentage of bipolar cells by counting cells that had both cell ends stained brightly as bipolar.

#### Orb6-as2 kinase inhibition

The design and construction of the *orb6-as2* analogue-sensitive mutant were as previously described^44^. Inhibition of Orb6-as2 kinase was achieved using the ATP analogue 1-NA-PP1 diluted in DMSO (5mM stock concentration) and stored at −20°C. In all experiments, the final concentration of 1-NA-PP1 used was 50 μM and cells were treated for 30 minutes.

#### CRISPR/Cas9 construction of Rga3 mutant strains

CRISPR/Cas9 was performed as previously described with some modifications^104^. We used CRISPR4P (<u>bahlerlab.info/crispr4p</u>) to select our sgRNA. We used the ligation-free method to clone our sgRNA into the CRISPR/Cas9 plasmid pMZ379. NEB® 10-beta Competent *E. coli* cells were transformed with our ligated plasmid. 12 colonies were selected, and the plasmids were sent for Sanger sequencing. As our HR template, we used single-stranded DNA containing our point mutation of interest as well as three (for Rga3) or four (for Wis1) silent point mutations between the point mutation of interest and the PAM site to prevent further Cas9 cleavage. Synchronized competent *S. pombe* cells were transformed using 20 µg denaturated salmon sperm DNA, 1 µg of HR template, 2 µg of sgRNA plasmid, and 145 μL 50% PEG4000. Cells were plated on YES plates with 100 μg/mL Nourseothricin. Once colonies were formed, 12-18 of the smallest colonies were streaked out onto YES plates without Nourseothricin to eliminate Cas9 plasmid. Colony PCR was performed and purified PCR samples were sent for sequencing to confirm successful point mutation construction.

#### Nitrogen starvation

All strains used in nitrogen starvation experiments were prototrophic and were cultured in unsupplemented EMM to eliminate the effect of supplemented amino acids as potential nitrogen sources. Therefore, the starvation of the nitrogen source in this study was achieved by removal of 0.5% NH_4_Cl. For starvation experiments, cells cultured in EMM+0.5% NH_4_Cl (EMM+N) were spun down, washed once in EMM+0% NH_4_Cl (EMMN), and resuspended in EMM+0% NH_4_Cl (EMM-N) for study.

#### Protein extraction and western blot analysis

Protein extraction was modified from previously described protocol^111^. ∼7.5 × 10^7^ total cells growing exponentially were harvested and cell pellets were resuspended in 300μL of H_2_O with protease and phosphatase inhibitor cocktail (Halt™ Protease and Phosphatase Inhibitor Cocktail (100X)) and transferred to 1.5mL microcentrifuge tube. 300μL of 0.6M NaOH with inhibitor cocktail (see above) was added to the cells and resuspended. Cells were left to incubate for 5 minutes at room temperature, inverting the tubes 2-3 times halfway through incubation. Cells were then centrifuged at 2300 RCF for 2 minutes. The supernatant was discarded, and the pellet was resuspended in 75μL modified SDS buffer (60mM Tris-HCl [pH 6.8], 4% 2-Mercaptoethanol, 4% SDS, 5% glycerol) with inhibitor cocktail (see above) and boiled at 98°C for 3 minutes. Cells were placed on ice and centrifuged at 4°C at 3500 RCF for 1 minute. 65μL of the supernatant (protein extract) was collected with 60μL stored at −80°C immediately, and 5μL used for downstream protein quantification assay. To quantify protein, the RC DC protein assay kit was used as it is compatible with the concentration of reducing agents and detergents present in the modified SDS buffer. Standard western blotting procedures were performed as follows: protein was separated using SDS-PAGE and transferred onto a nitrocellulose membrane. The membranes were probed with antibodies of interest and visualized through fluorescent detection on Li-Cor Odyssey 9120. Gef1-3YFP strains were used in detecting total Gef1 protein with anti-GFP and phosphorylated Gef1-S112 was detected using previously described antibody^41^. A ratio was determined between pGef1-S112 and total Gef1 unless otherwise specified. Quantification of the blots was performed using Image Studio software. Uncropped blots are provided in Fig. S4-6.

#### Stress-independent Sty1 activation

The stress-independent activation of Sty1 was performed as previously described^48^. Cells were grown in YES media at 30°C in the presence of 5μM 3BrB-PP1. 3BrB-PP1 was diluted in methanol for a stock solution concentration of 50mM and stored in −80°C. To activate Sty1, cells were washed twice in YES media with or without 5μM 3BrB-PP1 (without containing equal volume of methanol) and incubated with shaking for 30 or 60 minutes.

### QUANTIFICATION AND STATISTICAL ANALYSIS

Data are presented as described in legends. Experiments were completed in independent biological triplicates. A two-tailed unpaired Student’s t test was used to assess statistical significance between two groups. One-way or two-way analysis of variance (ANOVA) followed by appropriate post hoc test was applied to evaluate the difference between more than two groups (one-way) or more than 2 groups and more than one condition (two-way). Statistical analyses and visualization were performed with GraphPad Prism 10 (GraphPad Software, San Diego, CA). p-value < 0.05 was set as the threshold for statistical significance. Power analyses were performed using G*Power to determine minimum sample size^112,113^.

## SUPPLEMENTAL INFORMATION

**Video S1.** Active Cdc42 oscillatory dynamics, related to Figure 1A.

Fluorescent microscopy timelapse showing oscillations of active Cdc42 (CRIB-GFP) between cell tips.

**Video S2.** Active Cdc42 dynamics during Orb6 inhibition, related to Figure 1B.

Fluorescent microscopy timelapse showing exploratory pattern of active Cdc42 (CRIB-GFP) along the cell membrane when analogue-sensitive *orb6-as2* mutant is inhibited (+1-NA-PP1).

**Document S1.** Table S1-S2 and Figures S1-S6

