## Supplementary material for "Control of stress-activated Cdc42 dynamics by the MAP kinase Sty1 – NDR kinase Orb6 regulatory axis": DocumentS1

### Supplementary Figures

| Systematic ID | TSC | *orb6+* | *orb6ts* | *orb6ts/orb6+* | Protein | Description |
|---|---|---|---|---|---|---|
| SPAC8E11.02c | 7919 | 1000 | 1000 | 1.00 | Rad24 | 14-3-3 protein Rad24 |
| SPAC17A2.13c | 2250 | 179 | 449 | 2.51 | Rad25 | 14-3-3 protein Rad25 |
| SPAPB1E7.07 | 33 | 7 | 0 | 0.00 | Glt1 | glutamate synthase Glt1 (predicted) |
| SPAC17A2.14 | 77 | 16 | 1 | 0.04 | Spac17A2.14 | CorA family magnesium ion transmembrane transporter |
| SPCC4B3.04c | 34 | 7 | 0 | 0.05 | Nte1 | lysophospholipase (predicted) |
| SPAC6G10.02c | 101 | 20 | 1 | 0.05 | Tea3 | cell end marker Tea3 |
| SPBC9B6.04c | 32 | 6 | 0 | 0.05 | Tuf1 | mitochondrial translation elongation factor EF-Tu Tuf1 |
| SPAC17G6.15c | 30 | 6 | 0 | 0.05 | Spac17G6.15C | MTC tricarboxylate transmembrane transporter (predicted) |
| SPAC19A8.04 | 39 | 8 | 1 | 0.08 | Erg5 | C-22 sterol desaturase Erg5 |
| SPBC32F12.11 | 217 | 42 | 4 | 0.10 | Tdh1 | glyceraldehyde-3-phosphate dehydrogenase Tdh1 |
| SPAC21E11.08 | 31 | 6 | 1 | 0.11 | Lcb2 | serine palmitoyltransferase Lcb2 (predicted) |
| SPBC19C7.06 | 58 | 11 | 1 | 0.12 | Prs1 | cytoplasmic proline-tRNA ligase Prs1 (predicted) |
| SPBC27B12.11c | 30 | 6 | 1 | 0.17 | Pho7 | transcription factor Pho7 |
| SPBC354.12 | 135 | 25 | 5 | 0.20 | Gpd3 | glyceraldehyde 3-phosphate dehydrogenase Gpd3 |
| SPCC1223.08c | 78 | 14 | 3 | 0.20 | Dfr1 | dihydrofolate reductase/ serine hydrolase family fusion protein Dfr1 |
| SPBC1706.01 | 40 | 7 | 2 | 0.22 | Tea4 | tip elongation aberrant protein Tea4 |
| SPBC1A4.05 | 44 | 8 | 2 | 0.25 | Blt1 | ubiquitin domain-like protein Blt1 |
| SPBP35G2.07 | 64 | 11 | 3 | 0.26 | Ilv1 | acetolactate synthase catalytic subunit |
| SPBC32C12.03c | 41 | 7 | 2 | 0.27 | Ppk25 | serine/threonine protein kinase Ppk25 (predicted) |
| SPCC594.02c | 40 | 7 | 2 | 0.28 | Spcc594.02C | conserved fungal protein |
| SPBC56F2.12 | 33 | 6 | 2 | 0.28 | Ilv5 | acetohydroxyacid reductoisomerase (predicted) |
| SPCC970.08 | 86 | 14 | 5 | 0.36 | Spcc970.08 | inositol polyphosphate kinase (predicted) |
| SPBC1A4.02c | 32 | 5 | 2 | 0.36 | Leu1 | 3-isopropylmalate dehydrogenase Leu1 |
| SPAC4G8.13c | 47 | 8 | 3 | 0.42 | Prz1 | calcineurin responsive transcription factor Prz1 |
| SPAC12B10.14c | 31 | 5 | 2 | 0.46 | Tea5 | pseudokinase Tea5 |
| SPAC3H1.09c | 39 | 6 | 3 | 0.47 | Avt3 | vacuolar amino acid transmembrane transporter Avt3 |
| SPBC21B10.03c | 144 | 23 | 11 | 0.48 | Ath1 | ataxin-2 homolog |
| SPBP8B7.06 | 37 | 6 | 3 | 0.50 | Rpp201 | 60S acidic ribosomal protein A2 |
| SPAC29A4.11 | 125 | 19 | 10 | 0.52 | Rga3 | Rho-type GTPase activating protein Rga3 |
| SPAC1071.08 | 35 | 5 | 3 | 0.54 | Rpp203 | 60S acidic ribosomal protein A2 |
| SPAC9G1.06c | 379 | 58 | 32 | 0.55 | Cyk3 | cytokinesis protein Cyk3 (predicted) |
| SPCC16C4.09 | 141 | 21 | 13 | 0.62 | Sts5 | RNB-like protein |
| SPCC13B11.01 | 94 | 14 | 9 | 0.63 | Adh1 | alcohol dehydrogenase Adh1 |
| SPBC800.02 | 38 | 6 | 4 | 0.64 | Whi5 | cell cycle transcriptional repressor Whi5 (predicted) |
| SPBC16A3.07c | 31 | 5 | 3 | 0.64 | Nrm1 | MBF complex corepressor Nrm1 |
| SPAC24H6.09 | 72 | 11 | 7 | 0.65 | Gef1 | Cdc42 RhoGEF Gef1 |
| SPAC57A10.12c | 58 | 8 | 6 | 0.65 | Ura3 | dihydroorotate dehydrogenase Ura3 |
| SPCC297.03 | 182 | 26 | 18 | 0.68 | Ssp1 | Ca2+/calmodulin-dependent (CaMMK)-like protein kinase Ssp1 |

Table S1

**Supplementary Table 1: Mass spectrometry screen identifies potential Orb6 substrates.**

Mass spectrometry identified proteins were exported from Scaffold to Excel for further analysis. Contaminant proteins (e.g. keratins) were removed and nonspecific proteins that were also identified in TAP preparations from a strain lacking the TAP tag were shaded in gray. The spectral counts of the proteins from each purification are shown. For comparison, the spectral counts for each purification were first normalized to those of the bait – Rad24, and the ratio of the normalized spectral counts of "orb6ts/orb6+" was calculated. The ratio from lowest to highest is color coded as from blue to red, respectively. Only those proteins with the combined total spectral count of 30 or more and the ratio smaller than 0.7 are shown.

| Protein | Putative Orb6 phosphorylation consensus sequence (HX(R/K/H)XX(S/T) | Biological process |
|---|---|---|
| Blt1 | T177, S693 | Mitotic cytokinesis |
| Cyk3 | S300 | Mitotic cytokinesis |
| **Gef1** | **S112** | Cell polarity, mitotic cytokinesis |
| Tea3 | S430, S460 | Cell polarity |
| **Rga3** | S382, S419, **S683**, S751 | Cell polarity, signaling |
| Ppk25 | S38, S265, S404 | Signaling |
| Tea5 | S161 | Signaling |
| Kcs1 | S451 | Lipid metabolic process, kinase signaling |
| Nte1 | S1310 | Lipid metabolic process |
| Lcb2 | T265 | Lipid metabolic process |
| Dfr1 | T235 | Lipid metabolic process |
| **Sts5** | S84, **S86**, S169, S261, S638, S974 | mRNA metabolic process |
| Whi5 | S84, S86 | DNA-templated transcription |
| Nrm1 | S336 | DNA-templated transcription |
| Pho7 | S23, S429 | DNA-templated transcription, ion homeostasis |
| Prz1 | T79 | DNA-templated transcription, ion homeostasis |
| Mnr2 | S105 | Transmembrane transport, ion homeostasis |
| Fsf1 | S309 | Transmembrane transport, ion transporter |

Table S2

**Table S2: List of proteins that bind to Rad24 in an Orb6-dependent manner identified in TAP/MS screen with putative Orb6 phosphorylation consensus sequences.**

Tandem affinity purification (TAP) and mass spectrometry (MS) screen identified 29 Orb6 factors that bind to Rad24 in an Orb6-dependent manner. Shown in the table are the 18 proteins that contain the putative Orb6 phosphorylation consensus sequence [HX(R/K/H)XX(S/T)]. Previously identified Orb6 substrates Gef1 [S1] and Sts5 [S2,3] and their phosphorylation site are in bold, as well as Rga3, identified in this study. Sequence and GO biological process identified through PomBase.

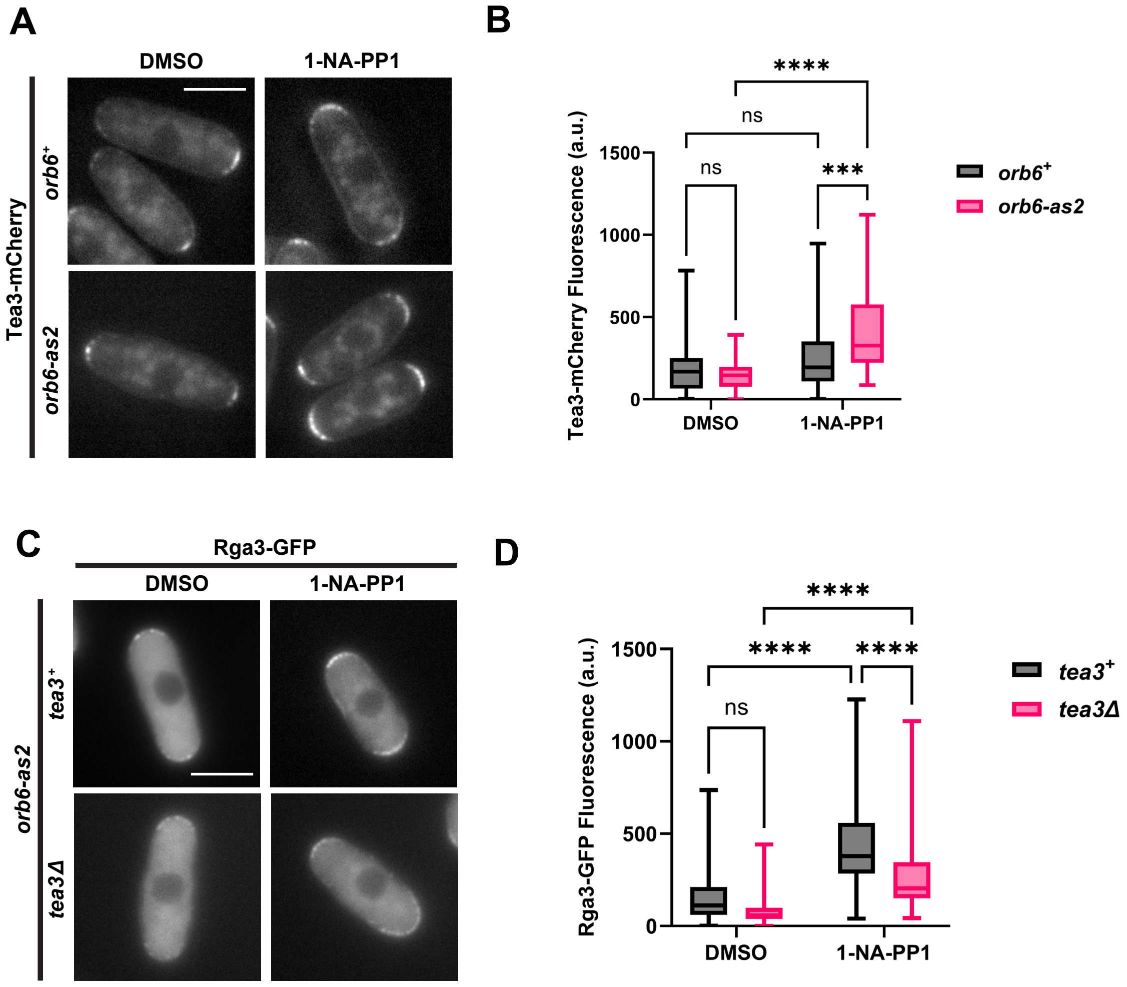
A
DMSO
1-NA-PP1
Tea3-mCherry
orb6+
orb6-as2
B
****
ns
ns
***
1500
1000
500
0
Tea3-mCherry Fluorescence (a.u.)
DMSO
1-NA-PP1
orb6+
orb6-as2
C
Rga3-GFP
DMSO
1-NA-PP1
orb6-as2
tea3+
tea3Δ
D
****
****
****
ns
1500
1000
500
0
Rga3-GFP Fluorescence (a.u.)
DMSO
1-NA-PP1
tea3+
tea3Δ

Figure S1

**Supplementary Figure 1: Tea3 localization increases upon Orb6 inhibition and promotes Rga3 localization to the cell tips.**

(A) Tea3-mCherry localization increases at the cell tips following Orb6 inhibition with 1-NA-PP1 for 30 minutes. Images are a sum projection Z-stack of 6 images separated by a step-size of 0.3 μm. Scale bar, 5 μm.

(B) Quantification of Tea3-mCherry localization at the cell tips depicted in (A) based on three independent experiments. Whiskers indicating minimum to maximum are shown, box represents $25^{th}$ to $75^{th}$ quartiles, and horizontal line represents median, *p* values are determined by two-way ANOVA with Tukey's HSD test, $p \leq 0.001$, ***, $p \leq 0.0001$, ****.

(C) Following Orb6 inhibition by 1-NA-PP1 for 30 minutes, Rga3-GFP localization decreases at the cell tips in the absence of Tea3. Images are a sum projection Z-stack of 6 images separated by a step-size of 0.3 μm. Scale bar, 5 μm.

(D) Quantification of Rga3-GFP localization at the cell tops depicted in (C) based on three independent experiments. Data presented as in (B), *p* values are determined by two-way ANOVA with Tukey's HSD test, $p \leq 0.0001$, ****.

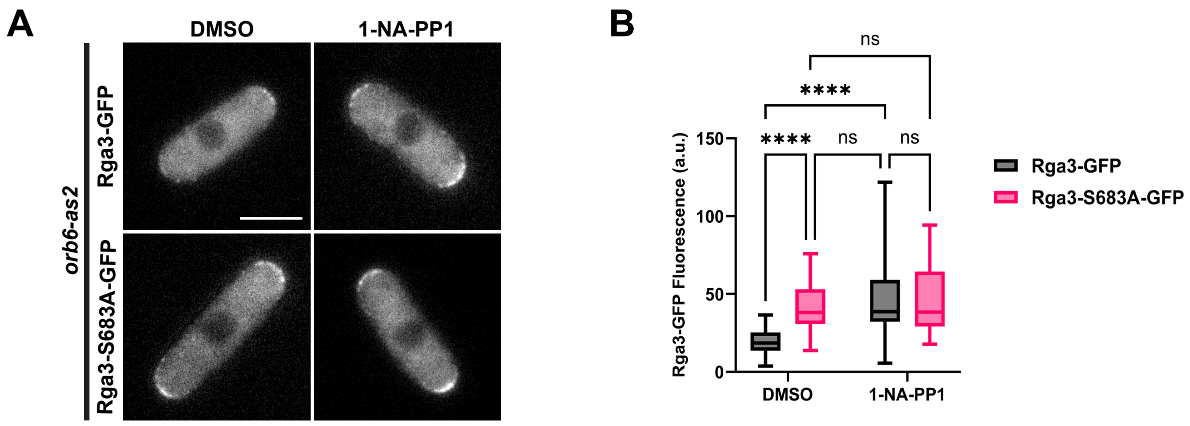
A
DMSO
1-NA-PP1
orb6-as2
Rga3-GFP
Rga3-S683A-GFP
B
ns
****
****
ns
ns
Rga3-GFP Fluorescence (a.u.)
150
100
50
0
DMSO
1-NA-PP1
Rga3-GFP
Rga3-S683A-GFP

Figure S2

**Supplementary Figure 2: Rga3-S683A localization is not further exacerbated by Orb6 inhibition.**

(A) Rga3-S683A-GFP localization does not change following Orb6 inhibition with 1-NA-PP1 for 30 minutes. Scale bar, 5 μm.

(B) Quantification of Rga3-GFP localization at the cell tips depicted in (A) based on three independent experiments. Whiskers indicating minimum to maximum are shown, box represents $25^{th}$ to $75^{th}$ quartiles, and horizontal line represents median, *p* values are determined by two-way ANOVA with Tukey's HSD test $p \leq 0.0001$, ****.

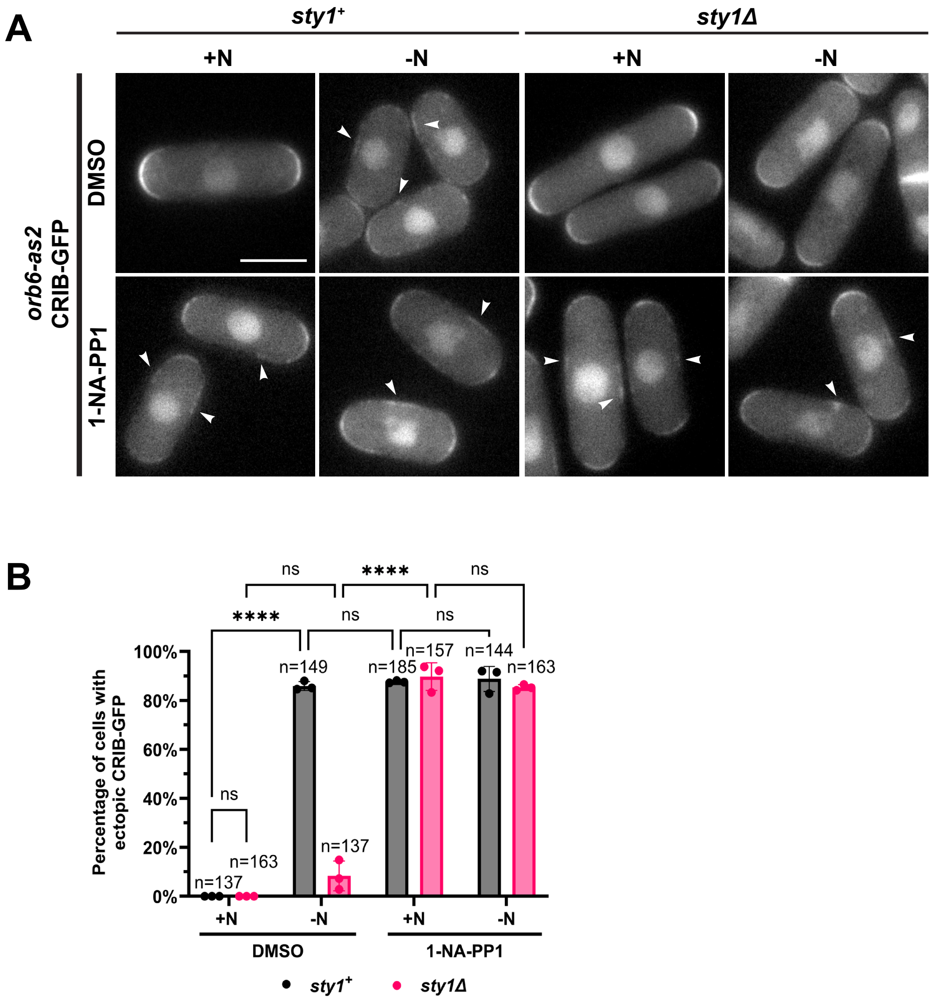
A
sty1+
sty1Δ
+N
-N
+N
-N
orb6-as2
CRIB-GFP
DMSO
1-NA-PP1
B
ns
****
ns
****
ns
ns
ns
Percentage of cells with ectopic CRIB-GFP
100%
80%
60%
40%
20%
0%
n=137
n=163
n=149
n=137
n=185
n=157
n=144
n=163
+N
-N
+N
-N
DMSO
1-NA-PP1
sty1+
sty1Δ

Figure S3

**Figure S3: Orb6 inhibition after nitrogen starvation induces exploratory Cdc42 dynamics in absence of *sty1*.**

(A) Active Cdc42 forms lateral patches at the membrane upon inhibition of *orb6-as2* mutants with 1-NA-PP1 for 30 minutes after exposure to nitrogen starvation 15 minutes prior. N represents nitrogen. Scale bar, 5 μm.

(B) Percentage of cells with ectopic CRIB-GFP localization from (A) quantification from (A) based on three independent experiments. Data are presented as mean ± SD, *p* values determined by two-way ANOVA with Tukey's HSD test $p \leq 0.0001$, ****. n = number of cells quantified.

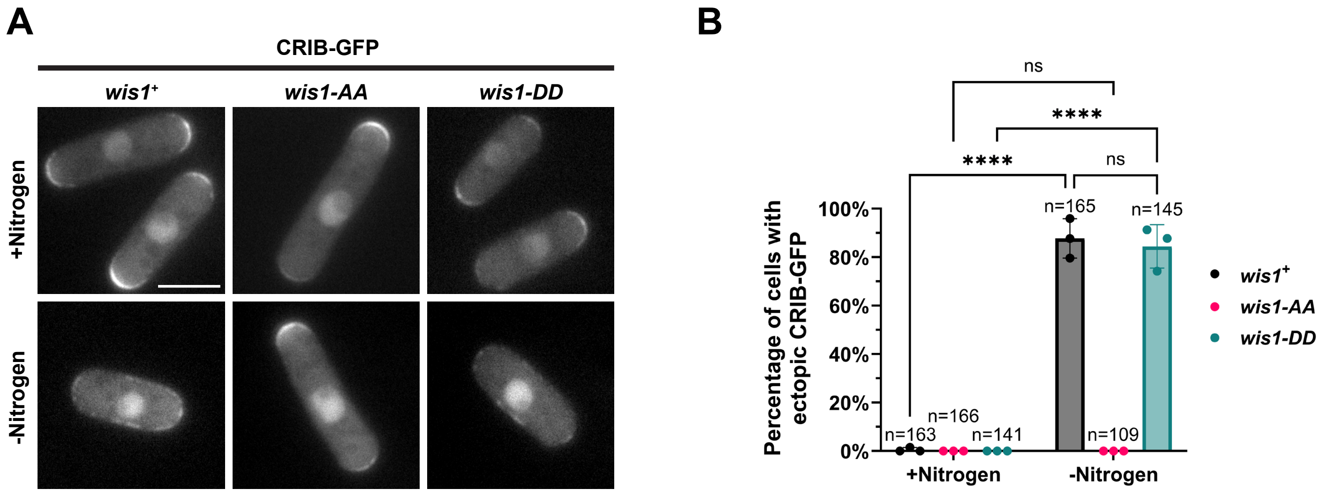
A
CRIB-GFP
wis1+
wis1-AA
wis1-DD
+Nitrogen
-Nitrogen
B
ns
****
****
ns
Percentage of cells with ectopic CRIB-GFP
100%
80%
60%
40%
20%
0%
n=163
n=166
n=141
n=165
n=109
n=145
+Nitrogen
-Nitrogen
wis1+
wis1-AA
wis1-DD

Figure S4

**Figure S4: Sty1 kinase activity promotes exploratory active Cdc42 dynamics during nitrogen starvation.**

(A) Active Cdc42 remains localized at the cell tips upon exposure to nitrogen starvation for 30 minutes in *wis1AA* mutants. Scale bar, 5 μm.

(B) Percentage of cells with ectopic CRIB-GFP localization from (A) quantification from (A) based on three independent experiments. Data are presented as mean ± SD, *p* values determined by two-way ANOVA with Tukey's HSD test $p \leq 0.0001$, ****. n = number of cells quantified.

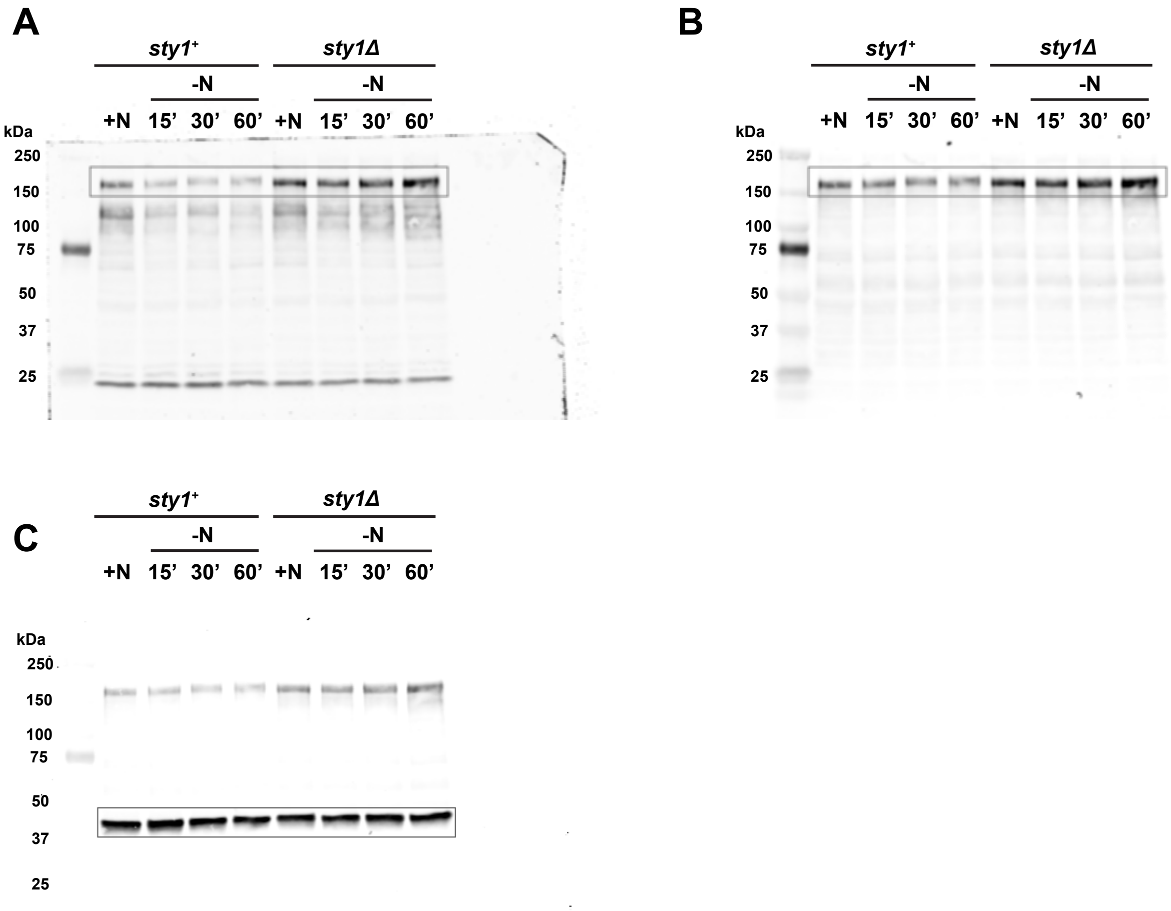
A
sty1+
sty1Δ
-N
-N
+N 15' 30' 60' +N 15' 30' 60'
kDa
250
150
100
75
50
37
25
B
sty1+
sty1Δ
-N
-N
+N 15' 30' 60' +N 15' 30' 60'
kDa
250
150
100
75
50
37
25
C
sty1+
sty1Δ
-N
-N
+N 15' 30' 60' +N 15' 30' 60'
kDa
250
150
100
75
50
37
25

Figure S5

**Supplementary Figure 5: Uncropped blot of Figure 5G.**

(A) Uncropped blot of pGef1-S112.

(B) Uncropped blot of tGef1.

(C) Uncropped blot of β-Actin.

A

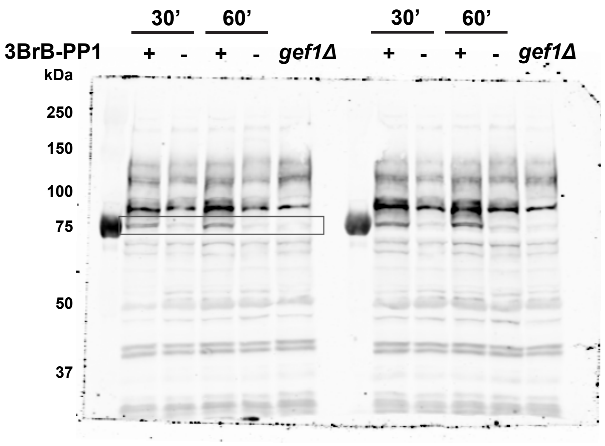
30'
60'
30'
60'
3BrB-PP1
+ - + - gef1Δ
+ - + - gef1Δ
kDa
250
150
100
75
50
37

B

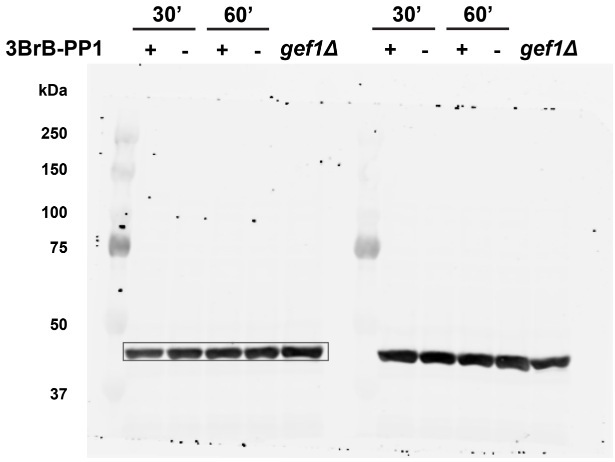
30'
60'
30'
60'
3BrB-PP1
+ - + - gef1Δ
+ - + - gef1Δ
kDa
250
150
100
75
50
37

Figure S6

**Supplementary Figure 6: Uncropped blot of Figure 5I.**

(A) Uncropped blot of pGef1-S112.

(B) Uncropped blot of β-Actin.

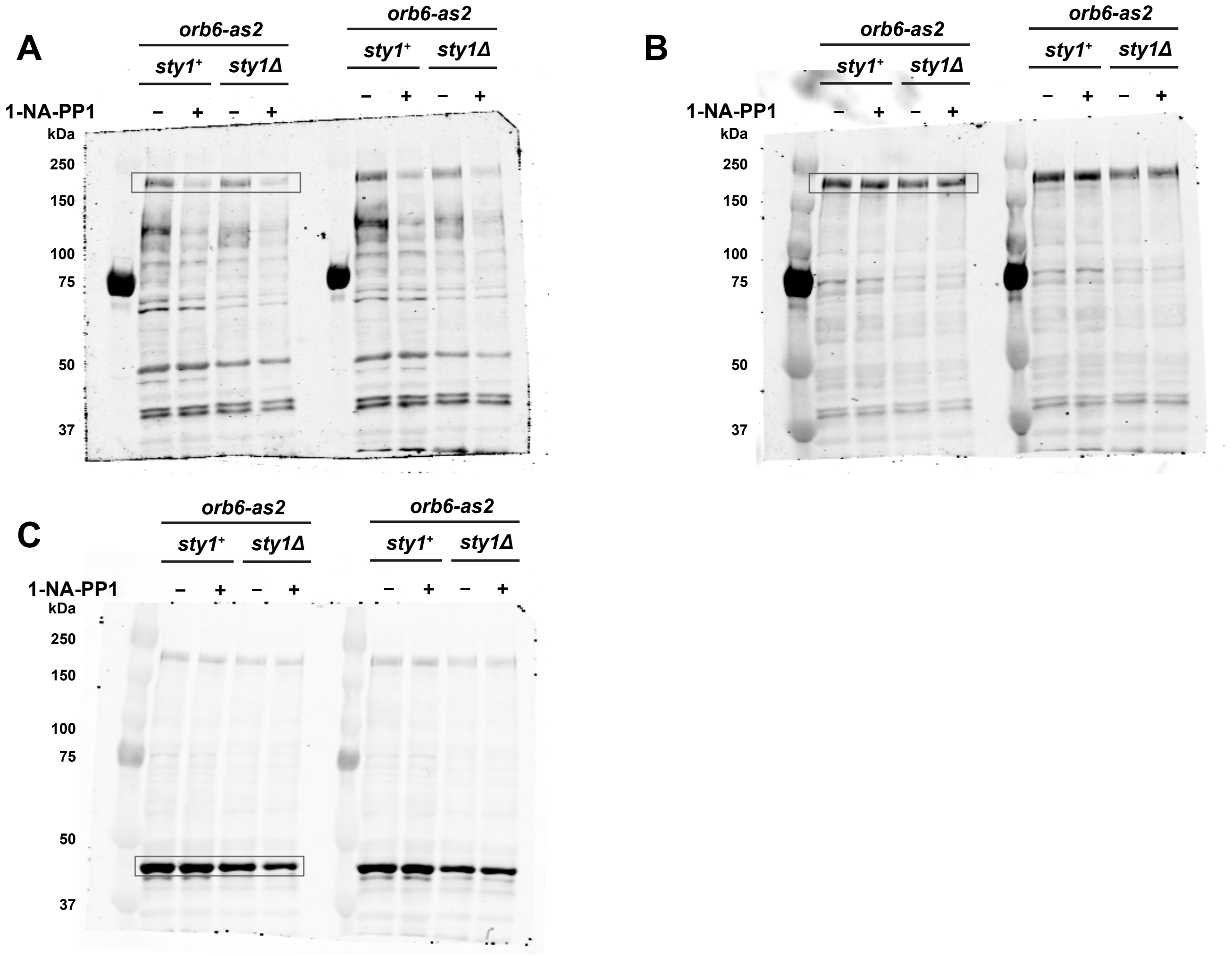


Figure S7

**Supplementary Figure 7: Uncropped blot of Figure 6A.**

(A) Uncropped blot of pGef1-S112.

(B) Uncropped blot of tGef1.

(C) Uncropped blot of β-Actin.
